# Learning from pre-pandemic data to forecast viral escape

**DOI:** 10.1101/2022.07.21.501023

**Authors:** Nicole N. Thadani, Sarah Gurev, Pascal Notin, Noor Youssef, Nathan J. Rollins, Chris Sander, Yarin Gal, Debora S. Marks

## Abstract

Effective pandemic preparedness relies on anticipating viral mutations that are able to evade host immune responses in order to facilitate vaccine and therapeutic design. However, current strategies for viral evolution prediction are not available early in a pandemic – experimental approaches require host polyclonal antibodies to test against and existing computational methods draw heavily from current strain prevalence to make reliable predictions of variants of concern. To address this, we developed EVEscape, a generalizable, modular framework that combines fitness predictions from a deep learning model of historical sequences with biophysical structural information. EVEscape quantifies the viral escape potential of mutations at scale and has the advantage of being applicable before surveillance sequencing, experimental scans, or 3D structures of antibody complexes are available. We demonstrate that EVEscape, trained on sequences available prior to 2020, is as accurate as high-throughput experimental scans at anticipating pandemic variation for SARS-CoV-2 and is generalizable to other viruses including Influenza, HIV, and understudied viruses with pandemic potential such as Lassa and Nipah. We provide continually updated escape scores for all current strains of SARS-CoV-2 and predict likely additional mutations to forecast emerging strains as a tool for ongoing vaccine development (evescape.org).

## Introduction

Viral diseases involve a complex interplay between immune detection in the host and viral evasion, often leading to the evolution of viral antigenic proteins. Antibody escape mutations affect viral reinfection rates and the duration of vaccine efficacy. Therefore, anticipating viral variants that avoid immune detection with sufficient lead time is key to developing optimal vaccines and therapeutics.

Ideally, we would be able to anticipate viral immune evasion by using experimental methods such as pseudovirus assays^1,2^ and higher-throughput deep mutational scans^3–17^ (DMSs) that measure the ability of viral variants to bind relevant antibodies. However, these experimental methods require antibodies or sera representative of the aggregate immune selection imposed on the virus, which only become available as large swaths of the population are infected or vaccinated, limiting the impact for *early* prediction of immune escape. In addition, since pandemic viruses can evolve rapidly (tens of thousands of new SARS-CoV-2 variants are currently sequenced each month), systematically testing all variants as they emerge is intractable, even without considering the effects of potential mutations on currently circulating strains.

It is therefore of interest to build computational methods for predicting viral escape that can be used to identify mutations that may emerge. An ideal model would be able to assess escape likelihood for as-yet-unseen variation throughout the full antigenic protein, would inform the design of targeted experiments, would be updated with pandemic information, and would make predictions with sufficient lead time for vaccine development (that is, before immune responses to the virus are observed). However, previous computational methods for forecasting viral fitness or immune escape depend critically on real-time sequencing or pandemic antibody structures, limiting their ability to predict unseen variants and making them impractical for vaccine development during the onset of a pandemic^18–22^.

In this work, we introduce EVEscape, a flexible framework that addresses the weaknesses of existing methods by combining a deep generative model trained on historical viral sequences with structural and biophysical constraints. Unlike existing methods, EVEscape does not rely on recent pandemic sequencing or antibodies, making it applicable both in the early stages of a viral outbreak and for ongoing evaluation of emerging SARS-CoV-2 strains. By leveraging functional constraints learned from past evolution, as successfully demonstrated for predicting clinical variant effects^23–25^, EVEscape can capture relevant epistasis^26,27^ and thereby predict mutant fitness within the context of any strain background. Moreover, EVEscape is adaptable to new viruses, as we demonstrate in both our validation on SARS-CoV-2, HIV, and Influenza and in predictions for the understudied Nipah and Lassa viruses. This approach enables advanced warning of concerning mutations, facilitating the development of more effective vaccines and therapeutics. Such an early warning system can guide public health decision-making and preparedness efforts, ultimately minimizing the human and economic impact of a pandemic.

## Results

### EVEscape combines deep learning models and biophysical constraints

Viral proteins that escape humoral immunity disrupt polyclonal antibody binding while retaining protein expression, protein folding, host receptor binding, and other properties necessary for viral infection and transmission^9^. We built a modeling framework—EVEscape—that incorporates constraints from these different aspects of viral protein function learned from different data sources. We express the probability of a mutation to induce immune escape as the product of three probabilities; the likelihood that a mutation maintains viral fitness (‘fitness’ term), occurs in an antibody accessible region (‘accessibility’ term), and disrupts antibody binding (‘dissimilarity’ term) (Figure 1A). These components are amenable to pre-pandemic data sources, allowing for early warning (Figure 1B).

**Figure 1:**
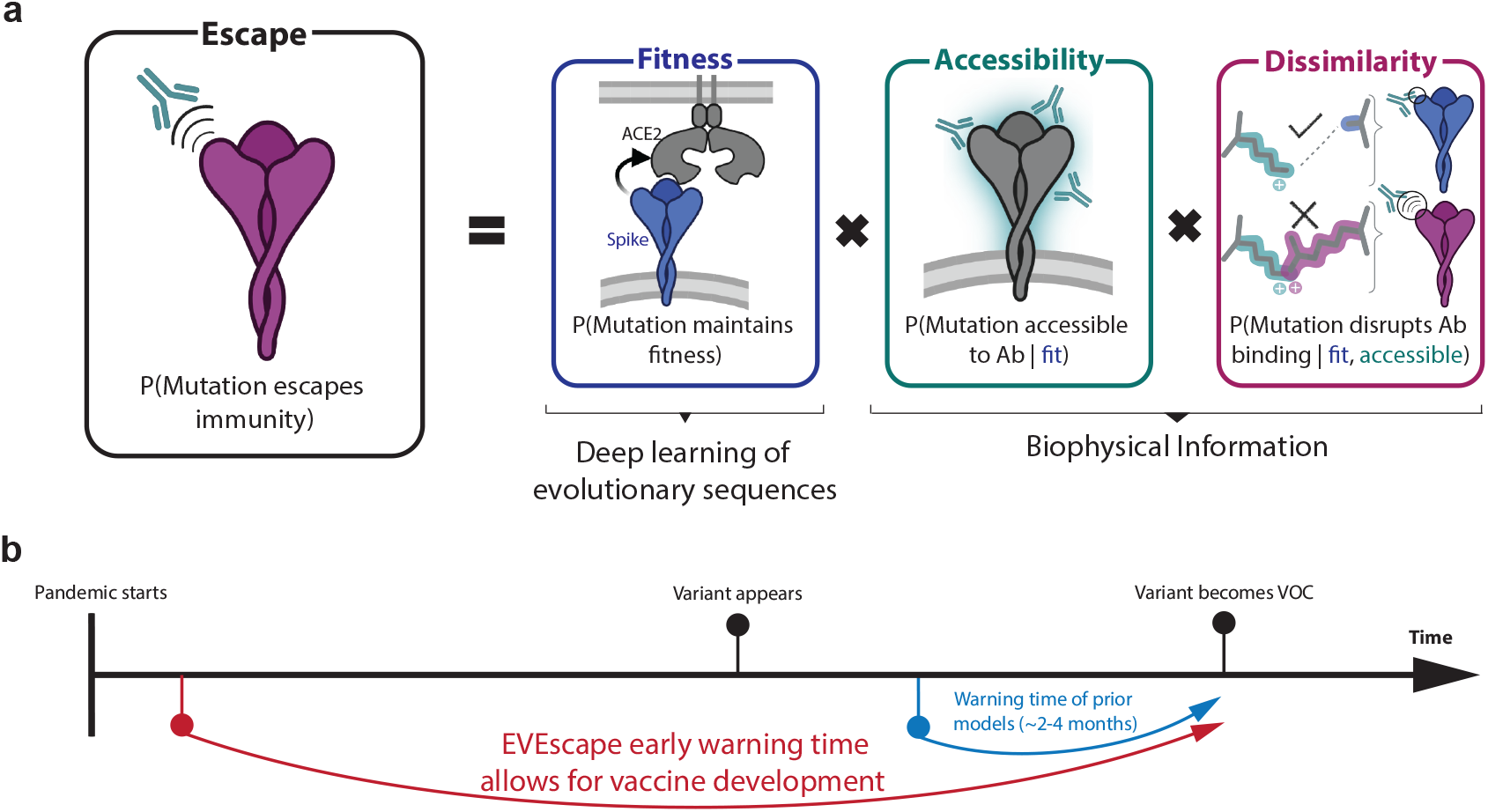
Early prediction of antibody escape from deep generative sequence models, structural and biophysical constraints. EVEscape assesses the likelihood of a mutation to escape the immune response based on the probabilities of a given mutation to maintain viral fitness, to occur in an antibody epitope, and to disrupt antibody binding. It only requires information available early in a pandemic, before surveillance sequencing, antibody-antigen structures or experimental mutational scans are broadly available.

Firstly, we estimate the fitness effect of substitution mutations (subsequently referred to as mutations) using EVE^23^, a deep variational autoencoder trained on evolutionarily-related protein sequences (Table S1-S2, Data S1) that learns constraints underpinning structure and function for a given protein family. Consequently, EVE considers dependencies across positions (epistasis), capturing the changing effects of mutations as the dominant strain backgrounds diversify from the initial sequence^28–30^. We showcase the efficacy of EVE by comparing model predictions and data from mutational scanning experiments that measure multiple facets of fitness for thousands of mutations to viral proteins^30–37^. Model performance approaches the correlation (ρ) between experimental replicates, including viral replication for influenza^31^ (ρ = 0.53) and HIV^30^ (ρ = 0.48) (Figure S1-S2, Data S2, Table S3). For SARS-CoV-2, we trained EVE across broad pre-pandemic coronavirus sequences, from sarbecoviruses like SARS-CoV-1 to “common cold” seasonal coronaviruses like the Alphacoronavirus NL63 (Table S1, Data S1), and compared predictions to measures of expression (ρ = 0.45) and receptor binding^35^ (ρ = 0.26) (Figure S1-S3). We note that sites which express in the DMS experiments but are predicted deleterious by EVE are frequently in contact with non-assayed domains of the Spike protein or with the trimer interface – interactions not captured in the RBD yeast-display experiment (Figure S3).

The second model component, antibody accessibility, is motivated by the need to identify potential antibody binding sites without prior knowledge of B cell epitopes. Accessibility of each residue is computed from its negative weighted residue-contact number across available 3D conformations (without antibodies), which captures both protrusion from the core structure and conformational flexibility^38–41^ (Figure S4, Table S4). Finally, dissimilarity is computed using differences in hydrophobicity and charge, properties known to impact protein-protein interactions^42–44^. This simple metric correlates with experimentally measured within-site escape more than individual chemical properties, BLOSUM substitution-matrix derived distance^45^, or distance in the latent space of the EVE model (Figure S5). To support modularity and interpretability of the impact of each component, each term is separately standardized and then fed into a temperature-scaled logistic function (Methods, Data S3-S4).

### Anticipating pandemic variation with pre-pandemic data

Extensive surveillance sequencing and experimentation prompted by the COVID-19 pandemic have presented a unique opportunity to assess EVEscape’s ability to predict immune evasion before escape mutations are observed^46,47^. To test the model’s capacity to make *early* predictions, we carried out a retrospective study using only information available *before* the pandemic (training on Spike sequences across *Coronaviridae* available prior to January 2020; Table S1, Data S1). We then evaluated the method by comparing predictions against what was subsequently learned about SARS-CoV-2 Spike immune interactions and immune escape.

The top predicted escape mutations for the whole of Spike are strongly biased towards the receptor-binding domain (RBD) and N-terminal domain (NTD), coincident with the bias for antigenic regions seen in the pandemic^48,49^ (Figures 2A-B, Figure S6). Within these domains, EVEscape scores are biased towards neutralizing regions—the receptor-binding motif of the RBD and the neutralizing supersite^50^ in the NTD (Figure 2C). EVEscape’s ability to identify the most immunogenic domains of viral proteins without knowledge of specific antibodies or their epitopes could provide crucial information for early development of subunit vaccines in an emerging pandemic^51^.

**Figure 2:**
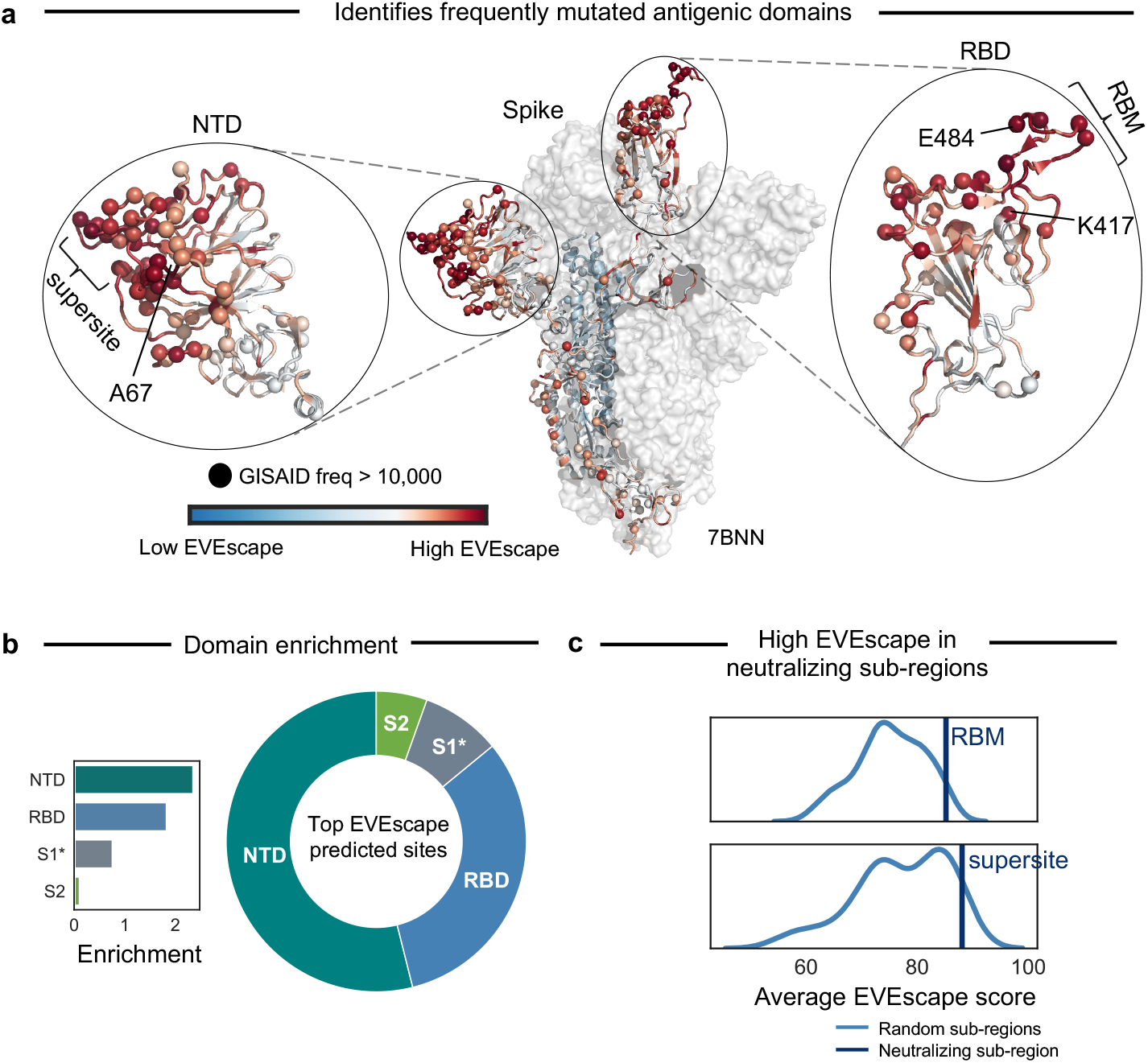
EVEscape identifies antigenic regions without antibody information. a) EVEscape scores mapped onto a representative Spike 3D structure (PDB identifier: 7BNN) highlight high-scoring regions with many observed pandemic variants, both in the RBD (receptor-binding domain) and NTD (N-terminal domain). Spheres indicate sites with mutations observed more than 10,000 times in the GISAID sequence database. b) The top decile of EVEscape predictions span diverse epitope regions across Spike, but the majority of predictions are in the NTD and RBD, which have a disproportionately high number of predicted EVEscape sites relative to their sequence length (enrichment). The regions considered are NTD (sequence positions 14 - 306), RBD (319 - 542), S1* (543 - 685), and S2 (686 - 1273), where S1* refers to the region in S1 between RBD and the S2. c) Neutralizing sub-regions – RBM (receptor-binding motif, 438-506) and NTD supersite^50^ (14-20,140-158, 245-263) – have significantly higher than average EVEscape scores, relative to a distribution of 150 random contiguous regions of the same length within the RBD and NTD, respectively.

We next compare model predictions to mutations that were subsequently observed in the pandemic as deposited in GISAID (Global Initiative on Sharing All Influenza Data)^46^, which contains over 500,000 unique sequences with over 12,000 missense mutations to Spike. For this analysis we focus on the RBD of Spike as this domain has been the most extensively studied due to its immunodominance^48,49^.

49% of our top RBD predictions were seen in the pandemic by December 2022 (Figure 3A, Figure S7; this proportion is robust to the threshold defining top escape mutations). The more often a mutation occurred in the pandemic, the more likely it is to be predicted by our method — 57% of high frequency observed substitutions are in the top EVEscape predictions (Figures 3B-C). We expect that the highest frequency mutations, seen in historical Variants of Concern (VOCs), will be enriched for escape variants that provide a fitness advantage in an immune population (whilst not expecting that all single substitutions in the VOCs will contribute to escape).

**Figure 3:**
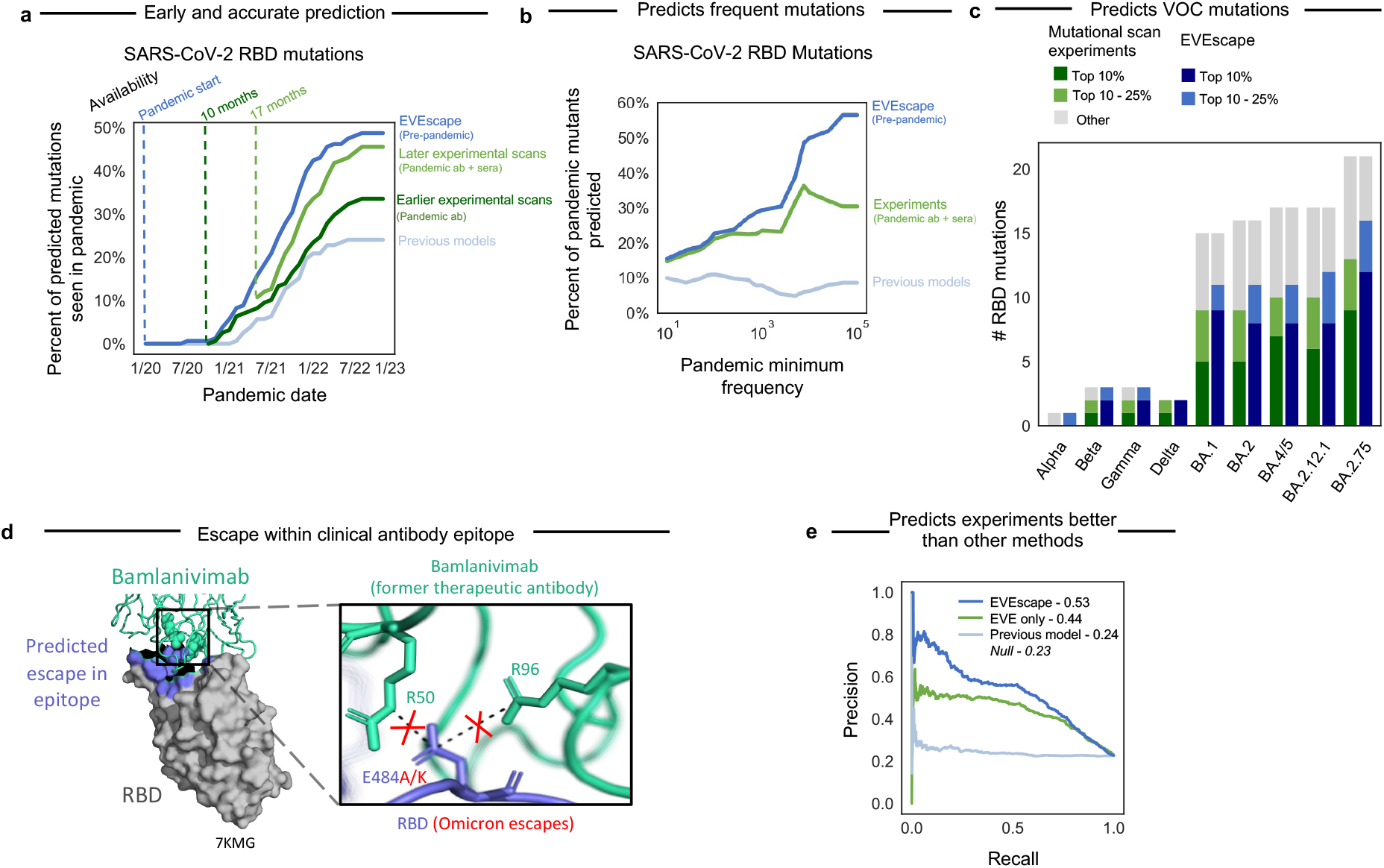
Pre-pandemic EVEscape is as accurate as intra-pandemic experimental scans at anticipating pandemic variation: retrospective analysis. a) Percent of top decile predicted escape mutations by EVEscape, mutational scan experiments (Bloom Set, Table S5), and a previous computational model^52^ seen over 100 times in GISAID by each date since the start of the pandemic. EVEscape based on pre-pandemic sequences anticipates pandemic variation at least on par with mutational scan experiments based on antibodies and sera available 10 or 17 months into the pandemic. Analysis focuses only on nonsynonymous point mutations that are a single nucleotide distance away from the Wuhan viral sequence. RBD is the receptor-binding domain of the Spike protein. b) Percent of observed pandemic mutations in top decile of escape predictions by observed frequency during the pandemic. High-frequency mutations in particular are well-captured by EVEscape. c) The majority of RBD mutations observed in VOC strains have high EVEscape scores and somewhat lower scores in the mutational scan experiments against pandemic sera. d) EVEscape can predict escape mutations in the epitope of the former therapeutic antibody bamlanivimab. E484 is involved in a salt bridge with R96 and R50 of bamlanivimab, which lost FDA Emergency Use Authorization due to Omicron’s emergence, wherein E484A or E484K mutations (both predicted in the top 1% of EVEscape Spike predictions) escape binding due to the loss of these salt bridges^53^. e) Precision-recall curve of RBD escape predictions of EVEscape, EVEscape fitness component only (EVE model) and previous computational model^52^ when compared to DMS escape mutations (AUPRC reported with a comparison to a “null” model where escape mutations are randomly predicted).

Not surprisingly, the fitness model component alone (here EVE^23^) is better that the full EVEscape model at predicting mutations seen at low frequency in the pandemic – likely because these mutations retain viral function but do not necessarily affect antibody binding or have a strong fitness advantage over other strains (Figures S7-S8). This suggests that EVEscape’s immune-specific components reflect important pandemic constraints and allow for mutation interpretability. For instance, VOC mutations R190S and R408S, with high EVEscape but low EVE scores, are in hydrophobic pockets that may facilitate significant immune escape^54^ (Figure S8). Meanwhile, the few VOC mutations (i.e., A222V and T547K) with significant EVE— but not EVEscape—scores have functional improvements such as monomer packing and RBD opening but do not impact escape^55,56^ (Figure S8). We also see that the proportion of EVEscape predictions seen during the pandemic increased over time—from 3% in December 2020 to 49% in December 2022 (Figure 3A)—and should continue to increase, an expected trend both as more variants are observed and as adaptive immune pressure increases^57^ with the growing vaccinated or previously infected population. Similarly, the fraction of mutations in VOC strains with high EVEscape scores has also increased (Figure S7).

Our model also predicted escape mutations that were subsequently observed in the pandemic in the epitopes of well-known therapeutic monoclonal antibodies under current or former Emergency Use Authorization^47^ (Figure S9), e.g., N440, E484A/K/Q, and Q493R. These predictions demonstrate the interplay of our three model components; for instance, the high accessibility as well as mutability of E484 results in 50% of all possible mutations at this site in the top 2% of EVEscape predictions and includes E484A/K in the top 1%—notable for escape from bamlanivimab^53^ (Figure 3D)—because of their high dissimilarity scores. We also identify candidate escape mutations in these therapeutic epitopes that have not yet been observed at frequencies higher than 10,000 – for instance variants to K444 and K417 (Figure S9), a subset of which are beginning to appear. This result suggests that escape sites can be well predicted before a pandemic and may have concrete applications for escape-resistant therapeutic design and early warning of waning effectiveness.

EVEscape represents a significant improvement over past computational methods. EVEscape is more than twice as predictive as prior unsupervised models^52^, both at predicting pandemic mutations (49% vs. 24% of top predictions observed in pandemic and 57% vs. 9% of highest frequency mutations predicted) as well as experimental measures of antibody escape (0.53 vs. 0.24 AUPRC) (Figures 3A-B, Figure 3E, Figure S7, Figures S10-S11, Figure S14, Table S5). All EVEscape components play a role in these predictions, with fitness predictions and accessibility metrics identifying sites of escape mutations while dissimilarity identifies amino acids that facilitate escape within sites (Figure S12-13). Moreover, other computational methods^20,22^ focus on near term prediction of strain dominance rather than longer term anticipation of immune evasion as they rely on pandemic sequences, antibody-bound Spike structures, or both, thereby hindering the ability to assess early predictive capacity. It is therefore notable that EVEscape outperforms even supervised approaches at predicting mutations seen in the pandemic (Figure S7).

### Comparative accuracy of EVEscape and high-throughput experiments

We contextualize the performance of EVEscape in comparison to deep mutational scans (DMS), which have been invaluable in identifying and predicting viral variants that may confer immune escape^3–13^. However, these experiments require polyclonal or monoclonal antibodies from infected or vaccinated people, limiting their early predictive capacity. For example, the DMS experiments conducted by 17 months into the pandemic (using 36 antibodies and 55 sera samples) are a third more predictive (46% vs. 34% observed) than the experiments conducted 7 months prior (using just 10 antibodies) (Figure 3A, Figure S7).

Despite being computed on sequences available more than 17 months earlier, EVEscape is as good as, or better than, the latest DMS scans at anticipating pandemic variation (49% vs. 46% observed, respectively, when considering the top decile of prediction) (Figure 3A). As we consider higher frequency mutations, EVEscape increasingly predicts a greater portion of pandemic variation than experiments (Figure 3B) and predicts a higher fraction of mutations in VOC strains (Figure 3C).

Discrepancies between EVEscape and experiments shed light on the complementary strengths of these approaches. EVEscape and experiments miss 41 and 46 pandemic mutations, respectively, that are predicted by the other method (Figure 4A, Figure 4D). These differences could indicate model inaccuracies or could reflect sparse sampling of host sera response in DMS experiments as well as artifacts from experiments testing only the RBD domain and missing the full set of *in vivo* constraints. Indeed, as more antibodies are incorporated in experiments, the agreement between EVEscape and experimental predictions increases (Figure S14). The majority of high EVEscape predictions that are not observed in experimental predictions are in known antibody epitopes (Figure 4B, Figure S13). By contrast, those mutations identified by the experiments that are below the threshold in EVEscape predictions are often predicted to have low fitness due to high conservation in the alignment at those positions.

**Figure 4:**
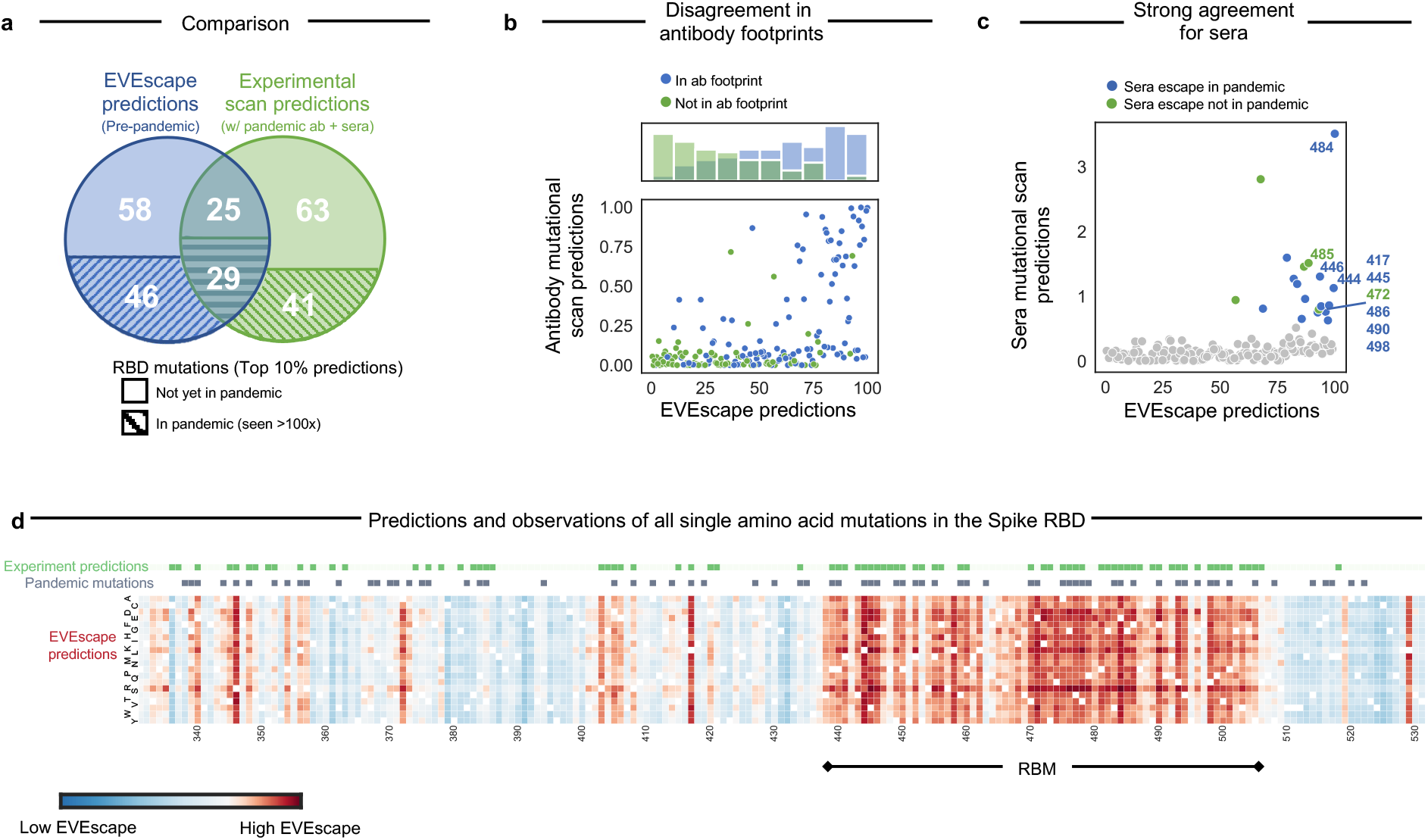
EVEscape and experiments make distinct, complementary escape predictions. a) Share of top decile of predicted escape mutations, predicted using EVEscape or based on mutational scan experiments (Bloom Set, Table S5), seen so far over 100 times in the pandemic. As the virus evolves further, more of the predicted escape mutations are expected to appear. b) RBD site-averaged EVEscape scores agree with site-averaged antibody escape experimental mutational scan measures (Bloom Set, Table S5), with high EVEscape sites that are missing from experimental escape prediction found within known antibody footprints. Hue indicates known antibody footprints from the PDB (information that EVEscape does not use as a pre-pandemic model). c) Predicted escape mutations based on experimental mutational scans (Bloom Set, Table S5) measuring recognition by convalescent sera from patients infected with either Wuhan, Beta, or Delta have high EVEscape scores. Mutations that escape sera are colored by whether they have occurred in the pandemic over 100 times. d) Heatmaps illustrating the EVEscape scores of all single mutations to the Wuhan sequence of SARS-CoV-2 RBD. Top lines are sites with observed pandemic mutation frequency >100 and sites in the top 15% of DMS experimental predictions based on mutational scan experiments. RBM is the receptor-binding motif.

The consensus between EVEscape and experiments is also of interest. We see that agreement is especially strong for polyclonal patient sera (Figure S14); in fact, half of the top 10% of EVEscape RBD sites are sera escape sites from experiments^5–7,14,15^ (Figure 4C). These mutants are of particular interest since they escape from the unique composition of antibodies produced by convalescent patients and are thus crucial to considerations of reinfection and vaccine design. For instance, E484, mutated in several VOCs, has the highest experimental sera binding and is the top EVEscape predicted site.

### Adapting EVEscape to reflect pandemic characteristics through its modular framework

The modular design of our framework facilitates its adaptability to the specific characteristics of a pandemic and to new data as it becomes available. To consider the effects of insertions and deletions on SARS-CoV-2 Spike immune escape^58^, we replace the EVE fitness component with TranceptEVE^59^ – a recently developed protein large language model which has previously demonstrated state-of-the-art performance for mutation effects prediction, including indels, which both prior computational models and high-throughput experiments have been unable to capture for SARS-CoV-2. When applied to the pandemic, this model captures the most frequent single insertion and deletion, both at site 144, each in the top decile of pandemic and random indel predictions (Figure S15). We also show that including glycosylation in the dissimilarity component for HIV Env, where glycans play an important role in immune escape^60–63^, improves model predictions of high-throughput experimental escape^17^ (Area under the precision recall curve raises 10% when including glycosylation for HIV; Figure S16). Additionally, we retrain EVE models with the addition of 11 million new sequences collected during the pandemic, which helps improve agreement with fitness DMS experiments by 20% (Figures S1, S17). This model captures epistatic shifts between Wuhan and BA.2, identifying changes in mutation fitness in the RBD and near BA.2 mutations and predicting positive epistatic shifts for known convergent omicron mutations and likely-epistatic wastewater mutations^64^ (Figure S18).

### Strain forecasting with EVEscape

A key application of an escape prediction framework is to identify circulating strains with high immune escape potential soon after their emergence, thus enabling the deployment of targeted vaccines and therapeutics before their spread. While the World Health Organization seeks to identify new high-risk variants as they arise, new strains are occurring at an increasing rate with now tens of thousands of novel SARS-CoV-2 strains each month, a scale infeasible for experimental risk assessment^65^. To create strain-level escape predictions, we aggregated EVEscape predictions across all individual Spike mutations in a strain. We evaluated EVEscape strain predictions for their alignment with experimental measures of strain immune evasion as well as their identification of known escape strains from pools of random sequences and from other strains observed at the same pandemic timepoint.

First, we see that pre-pandemic EVEscape-strain scores correlate well with experiments quantifying vaccinated sera neutralization of 21 strains^22^ (ρ = 0.80; Figure 5A, Data S5), better than an existing computational strain-scoring method (ρ = 0.77)^22^ even though that method uses 332 pandemic antibody-Spike structures for the prediction. Second, we show that EVEscape-strain scores for VOCs are consistently higher than random sequences at the same mutational depth, and in particular the Beta and later Omicron BA.2, BA.4, BA.2.12.1, BA.2.75, and XBB strains are in the top 1% of these generated sequences (Figure S19). EVEscape strain scores for these VOCs are also in the top 2% against sequences composed only of mutations already known to be favorable — those seen more than 100 times in GISAID, and even more strikingly, against combinations of mutations sampled from other VOCs (Figure S19).

**Figure 5:**
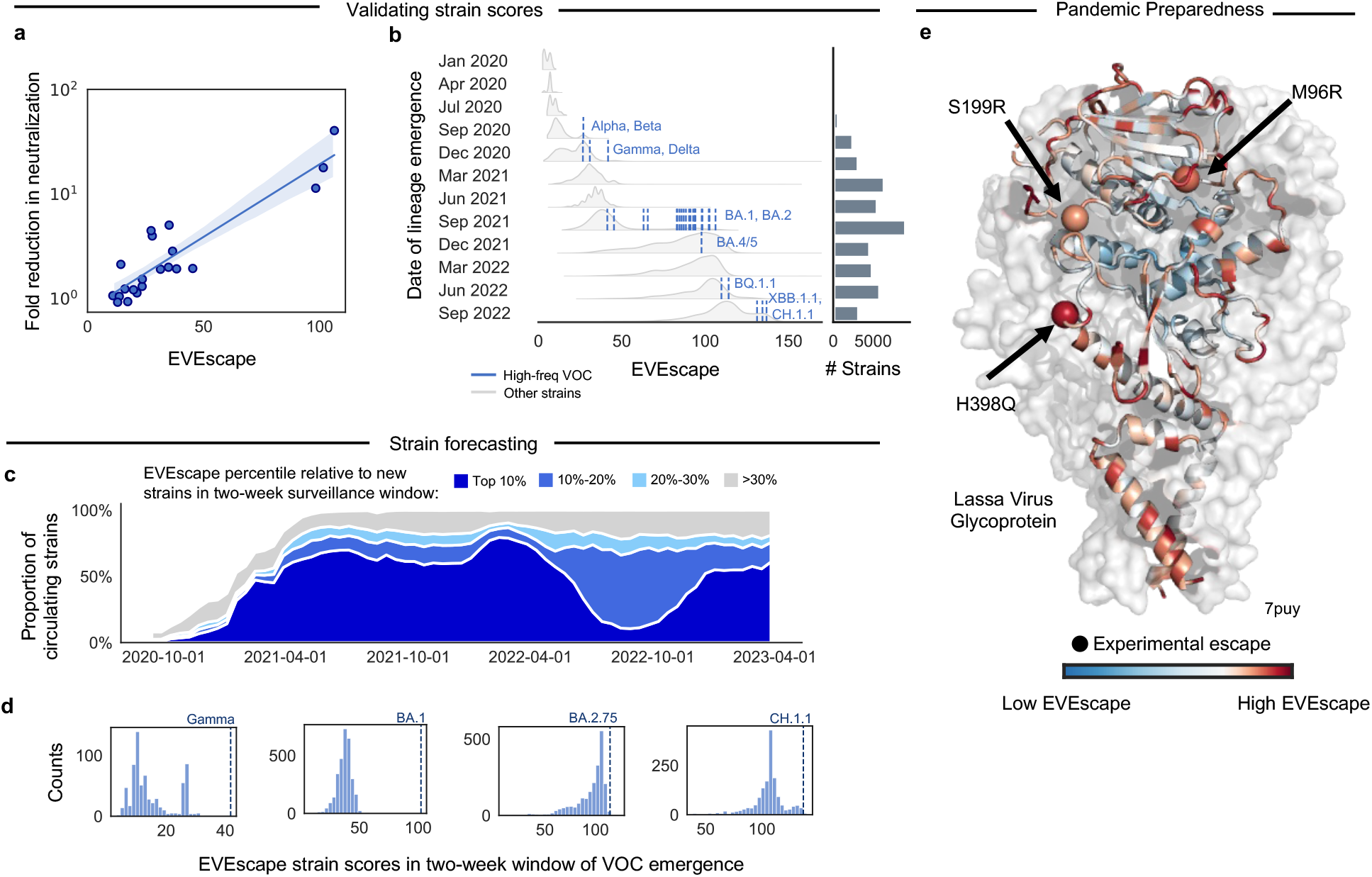
EVEscape applications: Identifying strains with high escape potential and forecasting escape for future pandemics. a) Pre-pandemic EVEscape scores computed for pandemic strains correlate with fold reduction in 50% pseudovirus neutralization titer^22^ for each strain relative to Wuhan (ρ = 0.80, n = 21). Linear regression line shown with a 95% confidence interval. b) Distributions of newly emerging EVEscape strain scores for non-VOCs (unique combinations of mutations) throughout 12 periods of the pandemic, with counts of the number of unique new strains per period. EVEscape strain scores increase throughout the pandemic. High frequency VOCs (occurring more than 5000 times) are shown in the first period each emerged, depicting that new VOCs are predicted to have higher escape scores than most strains in all previous time periods. c) Pandemic circulating strains are grouped according to their EVEscape decile relative to other strains emerging in the same non-overlapping two-week surveillance window. The relative prevalence of each EVEscape decile over the course of the pandemic is plotted in a stacked line-plot. The majority of circulating strains fall into the top 10% bin. Proportions do not sum to 100% as strains that emerged before the surveillance period of 9/2020 – 3/2023 are not included. d) VOCs (dotted lines) are among the highest scoring of hundreds or thousands of new strains (histograms) within their two-week window of emergence, enabling EVEscape to forecast which strains will dominate as soon as they appear after only a single observation. e) Site-wise maximum EVEscape scores on Lassa Virus Glycoprotein structure (PDB: 7PUY). We show agreement between sites of high EVEscape scores (in red) and escape mutations with experimental evidence (shown with spheres).

Lastly, we examine EVEscape’s ability to identify immune-evading strains as they emerged in the pandemic. We see that EVEscape scores have increased throughout the pandemic and that they are higher for more recent VOCs, reflecting their increased propensity for immune escape (Figure 5B). Moreover, EVEscape scores for newly emerging VOCs are higher than almost all strains in previous time periods (Figure 5B). Taken together, these results suggest EVEscape’s promise as an early-detection tool for picking out the most concerning variants from the large pool of available pandemic sequencing data. We therefore examine EVEscape’s utility as a tool to identify strains with high escape potential as they emerge in two-week surveillance windows. We see that the majority of circulating strains were in the top decile of EVEscape scores for their two-week window of emergence (Figure 5C). Moreover, in the two-week windows where the VOC strains Alpha, Beta, Gamma, Omicron BA.1, and Omicron BA.2.75 emerged, each VOC ranked in the top 5 of hundreds or thousands of new strains (Figure 5D, Figure S19). This demonstrates the ability of EVEscape to forecast which strains will dominate as soon as they appear after only a single observation, even as experimental testing of all emerging strains has become intractable.

To enable real-time variant escape tracking, we make monthly predictions (Data S5) available on our website (evescape.org), with EVEscape rankings of newly occurring variants from GISAID and interactive visualizations of likely future mutations to our top predicted strains. In sum, the EVEscape model captures relative immune evasion of successful strains and can identify concerning strains from pools of random combinations of mutations as well as from their temporal peers.

### EVEscape generalizes to other viral families with pandemic potential

Most viruses with pandemic potential have far less surveillance and research than SARS-CoV-2^66^. One of the main features of EVEscape is the ability to predict viral antibody escape before a pandemic—without the consequent increase in data during a pandemic—to narrow down vaccine sequences and therapeutics most likely to provide lasting protection, to assess strains as they arise, and to provide a watch list for mutations that might compromise any existing therapies. As one of the first comprehensive analyses of escape in these viruses, we applied the EVEscape methodology to predict escape mutations to the Lassa virus and Nipah virus surface proteins; these viruses cause sporadic outbreaks of Lassa hemorrhagic fever in West Africa and highly lethal Nipah virus infection outbreaks in Bangladesh, Malaysia, and India. Crucially, the three mutants present in the Lassa IV lineage that are known to escape neutralizing antibodies^67^ are all in the top 10% of EVEscape predictions, suggesting that EVEscape captures features relevant for Lassa glycoprotein antibody escape (Figure 5E). EVEscape predictions also identify 11 of 12 known escape mutants to Nipah antibodies^68–72^ (Figure S20).

Moreover, we demonstrate generalizability to Influenza Hemagglutinin^16^ and HIV Env^17^ based on DMS evaluation (Figure S10, Data S3). Based on these findings, we provide all single mutant escape predictions for these proteins (Data S5) to inform active and ongoing vaccine development efforts with the goal of mitigating future epidemic spread and morbidity.

## Discussion

One of the greatest obstacles for developing vaccines and therapeutics to contain a viral epidemic is the high genetic diversity derived from viral mutation and recombination, especially when under pressure from the host immune system. An early sense of potential escape mutations could inform vaccine and therapeutic design to better curb viral spread. Computational models can learn from the viral evolutionary record available at pandemic-onset and are widely extensible to mutations and their combinations. However, novel pandemic constraints (such as immunity) are unlikely to be captured. To achieve early escape prediction, EVEscape combines a model trained on historical viral evolution with a biologically informed strategy using only protein structure and biophysical constraints to anticipate the effects of immune selection. We demonstrate that EVEscape forecasts pandemic escape mutations and can predict which emerging strains have high escape potential through a retrospective analysis of the SARS-CoV-2 pandemic. This computational approach can preempt predictions from experiments that rely on pandemic antibodies and sera by many months while providing similar levels of accuracy.

EVEscape provides surprisingly accurate early predictions of prevalent escape mutations but cannot anticipate all constraints unique to a new pandemic to determine the precise trajectory of viral evolution. This method will be best leveraged in synergy with experiments developed to measure immune evasion and enhanced with pandemic data as it becomes available. Early in a pandemic, EVEscape can predict likely escape mutations for prioritized experimental screening with the first available sera samples – validated escape mutations could be strong candidates for multivalent vaccines. EVEscape can also identify structural regions with high escape potential, so therapeutic antibody candidates with few potential escape mutants in their binding footprint may be accelerated. Later in a pandemic, EVEscape can rank emerging strains, as well as mutants on top of prevalent strains, for their escape potential, flagging concerning variants early on for rapid experimental characterization and incorporation into vaccine boosters. The model can also be augmented to leverage current knowledge on virus-specific immune targeting and mutation tolerance from experimental and pandemic surveillance data. In return, our computational framework can inform this collective understanding by proposing escape variant libraries for focused experimental investigations.

EVEscape is a modular, scalable, and interpretable probabilistic framework designed to predict escape mutations early in a pandemic and to identify observed strains and their mutants that are most likely to thrive in a populace with widespread pre-existing immunity as the pandemic progresses. To this end, we provide EVEscape scores for all single mutation variants of SARS-CoV-2 Spike to Wuhan as well as scores for all observed strains and predictions of single mutation effects on the most concerning emerging strain backgrounds, with plans to continuously update with new strains. As the framework is generalizable across viruses, EVEscape can be used from the start for future pandemics as well as to better understand and prepare for emerging pathogens. To further accelerate broad and effective vaccine development, we provide EVEscape mutation predictions for all single mutations to Influenza, HIV, Lassa virus and Nipah virus surface proteins.

## Data and Code Availability

All data is provided in supplementary materials. Code is available on GitHub^[1]^.

## Acknowledgments

We thank members of the Marks laboratory and the OATML group for many valuable discussions. N.N.T. is supported by an NIH NIGMS F32 fellowship (GM141007-01A1). N.N.T. and N.J.R. were supported by the Chan Zuckerberg Initiative CZI2018-191853. S.G. is supported by a Takeda Fellowship. S.G. and N.Y. are supported by the Coalition for Epidemic Preparedness Innovations (CEPI). P.N. is supported by GSK and the UK Engineering and Physical Sciences Research Council (EPSRC ICASE award no. 18000077). Y.G. holds a Turing AI Fellowship (Phase 1) at the Alan Turing Institute, which is supported by EPSRC grant reference V030302/1. D.S.M. holds a Ben Barres Early Career Award by the Chan Zuckerberg Initiative as part of the Neurodegeneration Challenge Network, CZI2018-191853 and is supported by the Coalition for Epidemic Preparedness Innovations (CEPI). We acknowledge the authors and originating and submitting laboratories of the sequences from GISAID for sharing sequencing data (detailed acknowledgements in Data S6). Figure 1A created with BioRender.com.

## Additional Information

Supplementary Information is available for this paper. Correspondence and requests for materials should be addressed to Debora Marks.

Supplementary Information: This file contains supplementary figures S1-S20 and supplementary tables S1-S5.

Supplementary Data 1: Alignments used for EVE models for Lassa virus, Nipah virus, SARS-CoV-2, HIV, and Flu.

Supplementary Data 2: EVE and EVmutation scores for DMS fitness experiments for SARS-CoV-2, HIV, and Flu.

Supplementary Data 3: EVEscape scores for all SARS-CoV-2, HIV, Flu, Lassa virus, and Nipah virus, mutations. Includes pandemic counts and DMS escape experiment scores used for Spike.

Supplementary Data 4: EVEscape performance for selection of factor-specific temperature scaling.

Supplementary Data 5: EVEscape scores for all SARS-CoV-2 PANGO lineages and scores for strain neutralization variants.

Supplementary Data 6: Acknowledgements for all GISAID sequences.

## Methods

### Data acquisition

#### Training Data

##### Multiple sequence alignments for fitness models

For each viral protein, we construct multiple sequence alignments performing 5 iterations of the profile-HMM based homology search tool jackhmmer^73^ against the UniRef100 database^74^. As previously reported for EVE, DeepSequence, and EVcouplings, we generally keep sequences that align to at least 50% of the target sequence and columns with at least 70% coverage, except in the case of SARS-CoV-2 Spike where we use lower column coverage as needed (30-70%) to maximally cover experimental positions and significant pandemic sites^23–25^. For our pre-pandemic (pre-2020) alignment used as the primary model throughout this paper, we remove pandemic sequences using the “date of creation” variable from UniRef. We optimize search depth to maximize sequence coverage and the effective number of sequences (Neff) included after re-weighting similar protein sequences in the alignment within a Hamming distance cutoff (theta) of 0.01. To select sequence depth, we prioritized alignments with coverage >0.7L and Neff/L>1, or if this was not attainable, relaxed the requirements for Neff/L (Table S2).

##### Alignments with pandemic sequences

We construct an “evolutionary alignment” with non-SARS-CoV-2 sequences as described above using jackhmmer (with at least 50% sequence coverage, at least 30% column coverage, and theta of 0.01). We extract the full sequences pulled into the jackhammer alignment and re-align the sequences using super5^75^, then remove gapped positions relative to the Wuhan sequence. We also construct a “pandemic alignment” with all unique Spike sequences (with count >100) seen up until 11/27/21 (when BA.2 first appeared in GISAID). We then concatenate that “pandemic alignment” with the “evolutionary alignment” to create the final alignment.

##### Protein structures for accessibility calculation

For each viral surface protein, we selected crystal structures representing known structural states available to B-cell and antibody interactions (extracellular conformations) (Table S4). All heteroatoms and protein chains not part of the multimeric viral surface protein were removed.

#### Evaluation data

##### Antibody footprints

To identify known antibody footprints of viral surface proteins in the RCSB PDB^76^, we queried the database with the protein name and the word “antibody” and required that the source organism contain both “Homo sapiens” and the given virus name. Then for each structure we identified antibody and viral protein polymer entities and computed the antibody footprint as any residue with any atom within 3.5 angstroms of the antibody. Finally, we mapped footprints to the target viral protein sequence by using SIFTS to renumber all hits according to a UniProt ID, then used a MUSCLE multiple sequence alignment of the different UniProt sequences to map those hits to the target viral protein sequence. We use this same method to identify antibody footprints for specific clinical antibodies. For experimental evidence of clinical antibody escape susceptibility, we used the Stanford Coronavirus Antiviral & Resistance Database (CoV-RDB) susceptibility summary for monoclonal antibodies under emergency use authorization.^47^.

##### Deep mutational scans

We benchmark our models on a series of viral protein deep mutational scans^3–17,30–37^ (Table S3, Table S5). For each viral mutational scan, we select the variable or variables of protein fitness or antibody escape treated as primary in the publications. For mutants where the result is provided as residue frequencies observed at a given site (such as results expressed as preferences and processed by dms_tools2), we normalize the data at each site by dividing by the value of the wild-type residue. For the HIV analysis, we exclude antibody VRC34.01 due to its large spread of escape mutation distal to the epitope^77^. For SARS-CoV-2 RBD, we use only antibodies/sera escape data from the Wuhan sequence for our primary results. We also utilize data provided about the antibodies tested for the SARS-CoV-2 escape DMS studies, including the class of each antibody as well as the SARS-CoV-2 neutralization potency and sarbecovirus binding breadth^9^. We use the RBD dimeric ACE2 binding and expression DMS data for analysis^35^.

##### Pandemic sequencing data

We downloaded data on Spike variants and their deposit dates in the Global Initiative on Sharing All Influenza Data (GISAID) EpiCoV project database (www.gisaid.org)^46^ on 10/24/22. We further processed this data to get counts of combinations of mutations, the date of emergence, and PANGO lineage, as well as to get the month of emergence for each single mutation in Spike. We also downloaded consensus mutations for each PANGO lineage on 10/31/22 and mutation frequencies on 10/26/22 from Covid-19 CG^78^.

##### Lassa virus and Nipah virus antibody escape data

We aggregated data on single mutations resulting in escape from known Lassa and Nipah virus antibodies from literature studies with experimentally determined reduction in antibody binding, reduction in antibody neutralization, or emergence in growth selection experiments^67–72^.

##### Epistasis mutation sets

Our convergent omicron mutation set is defining mutations in Omicron lineages at sites 346, 444, 452, 460, and 486. This set is: L452R, N460K, F486V, K444N, L452M, F486I, R346T, F490S, K444M, K444T.

Our wastewater mutation set is the set of mutations from Smyth et al.^64^, which are mutations that were frequent in wastewater, but had rarely been seen clinically (pre-Omicron, mid 2021), so may be likely epistatic. This set is: Q493K, Q498Y, Q498H, T572N, H519N, H519Q.

##### Strain Neutralization data

We download neutralization data from Beguir et al.^22^, which contains the observed 50% pseudovirus neutralization titer (pVNT_50_) for 21 SARS-CoV-2 S protein variants. The pVNT_50_ reduction is relative to Wuhan. Neutralization is measured for n ≥ 12 sera collected after primary 2-dose vaccination by the Pfizer BioNTech vaccine (BNT162b2) and assessed against vesicular stomatitis virus (VSV)-based pseudoviruses with each S protein variant.

### Modeling approach

#### Overarching framework

We express the probability of a single amino acid substitution to lead to immune escape as the product of three conditional probabilities (Figure 1A):

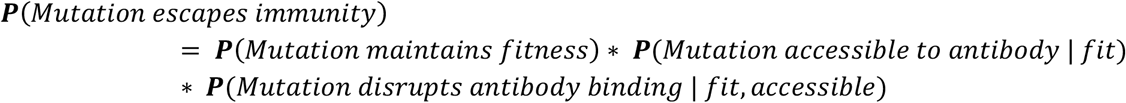

The EVEscape index estimates the log likelihood of escape as per the above equation. The fitness factor is obtained via a deep generative model for fitness prediction, while the accessibility and dissimilarity factors are features derived respectively from the known 3D structures for the viral protein and chemical characteristics of the amino acids involved in the mutation compared to the wild-type (see below for details).

Once selected, each factor is standardized and fed into a temperature scaled logistic function:

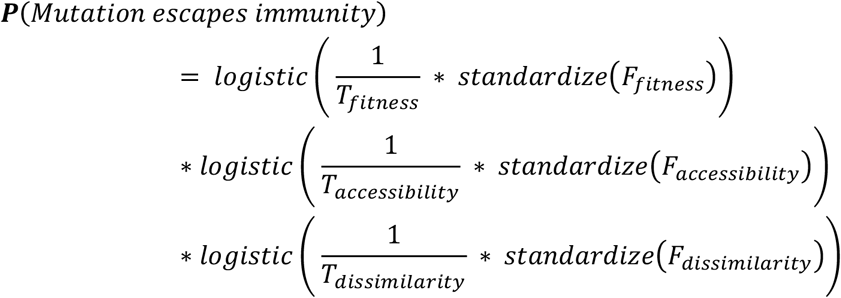

where the standardize(.) operator corresponds to standard scaling. We then take the log transform of the product of the 3 terms to obtain the final EVEscape scores.

Factor-specific temperature scaling helps recalibrate probability estimates for each term. We provide our hyperparameter grid search of these temperature hyperparameters across viruses in Data S4, examining versions of the model where we either include or do not include glycosylation in the dissimilarity term. We find that the fitness and accessibility components are already properly calibrated (*T*_*fitness*_ = *T*_*accessibility*_ = 1.0), while the dissimilarity component benefits from being slightly rescaled (*T*_*dissimilarity*_ = 2.0).

#### Fitness metric

Observed viral protein sequences reflect evolution under selection constraints for functional and infectious viruses. Generative sequence models express the probability that a sequence *x* would be generated by this process as *p*(*x*|*θ*), where the parameters *θ* capture the constraints describing functional variants. A generative model trained on observed viral protein variants can then be used to estimate the relative plausibility of a given mutant sequence as compared to wild-type by using the log ratio of sequence likelihoods as a heuristic:

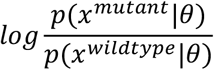

EVE (Evolutionary model of Variant Effects)^23^ is a Bayesian variational autoencoder (VAE)^79^, capable of capturing complex higher-order interactions across sequence positions. The fitness of a given protein sequence is measured via the log likelihood ratio of the mutated sequence *x* over that of the reference wild-type sequence *w*. Since an exact computation of the log likelihood of a sequence is intractable, we approximate it with the Evidence Lower Bound (ELBO) loss used to optimize the VAE:

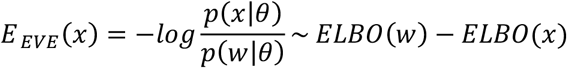

The ELBO term itself is estimated via Monte Carlo sampling, using 20k samples from the approximate posterior distribution. These approximations have been shown to provide strong results in practice^23^. Results are obtained by ensembling scores from 5 independently trained EVE models with different random seeds.

We train the different models following the procedure from the original EVE paper (see Frazer et al.^23^, Supplementary Section 3.2), using similarly-sized EVE models and with the same training hyperparameters. The only difference in our training procedure is that we slightly relax the constraint on minimum column coverage for sequences in the training MSAs (50% instead of 70%) as it led to superior fitness prediction performance in our hyperparameter tuning analyses for the different viruses modeled in this work.

In experiments aimed at illustrating the modularity of the EVEscape framework we leverage TranceptEVE, a recently developed protein language model with state-of-the-art performance for mutation effects predictions^59^. TranceptEVE is itself based off of two key components: 1) Tranception^80^, a family-agnostic autoregressive transformer trained on a large quantity of unaligned protein sequences from Uniref100^74^ from February 2022. 2) A family-specific EVE model that is trained to score sequences for a family of interest, and which acts as a prior distribution over amino acids at each sequence position. The predicted fitness for a given sequence is then obtained as a weighted average of the log likelihood assigned by these two components – the weights depending on the depth of the alignment used to train the underlying EVE model (deeper alignments implying a larger weight assigned to the EVE log likelihood).

For the experiments conducted in this work, we use the same ensemble of 5 EVE models as described above, as well as the large Tranception model checkpoint (∼700M model parameters) made available in Notin et al.^80^ which was trained on Uniref100 (see details of the training procedure in the corresponding paper in Appendix B.3).

#### Accessibility metric

Surface accessibility plays a key role in identifying where antibodies are most likely to contact a protein. While relative solvent accessibility (RSA) and weighted contact number (WCN) both reflect features of accessibility, we selected WCN as this metric also captures protrusion from the core structure that corresponds with where antibodies are known to bind proteins^38–41^ (Figure S4).

##### Calculating weighted contact number

We computed weighted contact numbers^41^ for each residue from structure as the sum of the square of the reciprocal distance between residue i and all other residues in the full protein (i.e., the full Spike trimer for SARS-CoV-2):

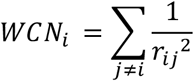

where *r*_*ij*_ is the distance between the geometric centers of the residue i and residue j side chains. Weighted contact number, beyond capturing surface accessibility, captures protrusion from the core structure and conformational flexibility^38–41^. By using squared distance, this value focuses on the degree of local interaction, and acts as a measure of exposure to the local environment that would permit antibody binding. It is both a simple and fast metric. We impute missing values in WCN due to gaps in the protein structure using the mean of WCN values of the residues preceding and following the gap.

##### RSA

We also explored RSA as a potential accessibility metric. To do so, we first computed accessible surface area based on hypothetical exposure to solvent water molecules using DSSP^81^. To calculate relative accessible surface area (RSA), we divided accessible surface area by the residue maximum accessibilities determined in Sander et al^82^. We impute missing values in RSA due to gaps in the protein structure by using the mean of RSA values of the residues preceding and following the gap (counting residues adjacent to the gap with RSA values>1 as part of the gap).

##### Aggregating across structures

When computing antibody-binding likelihood metrics across different structural conformations (i.e., both open and closed structures for SARS-CoV-2 Spike) we used the maximum accessibility (or minimum weighted contact numbers).

#### Dissimilarity metric

To predict the likelihood of a given mutation displacing an antibody interaction, we used a charge-hydrophobicity based measure of functional dissimilarity between the wild-type residue and the mutation residue. These are chosen as properties known to impact protein-protein interactions^42,43^. We compare our metric to individual chemical properties, substitution matrices, and the distance in the latent space of a VAE. We also experiment with incorporating glycosylation in our dissimilarity metric.

##### Charge-hydrophobicity

To compute a combined charge-hydrophobicity dissimilarity index, we standard-scaled the charge and hydrophobicity differences and then took the sum of the scaled differences. We use the Eisenberg-weiss hydrophobicity consensus scale^44^ and amino acid charge (as 1/0/-1) at physiological pH.

##### Chemical properties

We compared our metric to differences in residue size (side-chain mass), hydrophobicity, and charge.

##### Substitution Matrices

We compared our metric to the BLOSUM62^45^ matrix after dropping the null transition diagonal values.

##### Latent space distances

We also compared our metric to a metric of mutation distance learned by the EVE variational autoencoder. We calculated the L1 distance between the encoded representations of the wild-type viral protein sequence and a given single-mutation sequence in the latent space of the model, inspired by a similar approach first introduced by Hie et al.^52^

##### Glycosylation

We developed a version of our model considering glycosylation loss as a contributor to dissimilarity. While addition of glycosylation is also important for escape^60–63^, we focus here on loss of glycosylation for simplicity. In this version, we maximize the charge-hydrophobicity dissimilarity term if a mutation is likely to result in loss of a surface N-glycan site. We identified surface N-glycan sites as NxS/T sequons (where x is any amino acid except proline) with the N residue having an RSA>0.2. We consider that a mutation is likely to result in loss of glycosylation if the N or S/T is lost. We note that this can be an important factor for real-world escape even when some DMS experiments do not reflect the escape impacts of glycosylation loss, as is the case for SARS-CoV-2 experiments that use yeast display, with glycans different than in mammalian cells^4^. For HIV on the other hand, a significant portion of escape mutations from DMS experiments are a result of escape effects of glycan gains and loss^17^.

#### Imputing missing data

We impute missing values of features in EVEscape using the mean value of the feature across the target protein.

#### Insertions and Deletions

Scores for indels utilize tranceptEVE as the fitness component, negative weighted contact number as the accessibility component, and a maximized dissimilarity component score.

#### Strain-level EVEscape predictions

We aggregate across combinations of mutations by summing the EVEscape scores for each mutation.

### Evaluation

#### Comparison to functional assays

We compared model predictions to continuous experimental metrics of viral function using spearman’s rank correlation coefficient as our main evaluation metric, as previously described^24,25^.

#### Comparison to escape DMS

##### Data processing

As escape data is noisy at levels of low escape and a relatively low fraction of mutants exhibit escape, we chose to treat the escape outcome variable as binary. We selected a threshold for escape by fitting a gamma distribution to the data (combined across all screened antibodies and sera) and selecting the threshold corresponding to a 5% false discovery rate^17^. As the number of antibodies tested for RBD is much higher than for Flu and HIV, we bootstrapped the RBD data selecting 8 antibodies 1000 times and fitting a gamma distribution to these samples, then selected the average 5% false discovery rate threshold. As these thresholds are subject to our choice of a false discovery rate, we also plot performance for a range of thresholds (Figure S11). We identified a mutant as “escape” if its maximum escape value across any antibody tested exceeded the threshold — so a mutation for RBD is “escape” if it exceeds the threshold for any antibodies/sera in the Bloom or the Xie datasets (Data S3). We use thresholds of 0.57 for Bloom RBD, 0.9 for Xie RBD, 0.054 for Flu, and 0.138 for HIV to make model comparisons; mutations designated as escape by these experimental thresholds are almost all within 5Å of the antibody they escape (Figure S11). Note that the downloaded RBD escape datasets were already filtered using thresholds on expression and ACE2 binding of -1 and -2.35, respectively^83^.

To define a site-wise escape value, we averaged across the maximum escape values for each mutant at the site. For the antibody RBD DMS data, we define the antibody class of each mutation/site by determining the maximum number of antibodies for a given class that escape that mutation/site (Data S3).

As the scales are different for the Bloom and Xie datasets, we focus on the original Bloom RBD DMS data when we need to consider the top fraction of escape mutations. We examine performance on Flu and HIV as a secondary analysis to confirm generalizability, as fewer antibodies have been tested and the distribution of these antibodies does not reflect known immunodominant domains.

##### Metrics

To compare computational model performance in classifying escape mutants, we computed two metrics. We consider area under the receiver operating curve (AUROC) and area under the precision-recall curve (AUPRC). A key feature of an escape mutant predictor is the quality of its positive ‘escape’ predictions, as in practice, the positive predictive value will influence costly experimental screening efforts and selection of a limited number of variants for vaccine incorporation. To reflect this, we focus on the area under the precision-recall curve (AUPRC) as a performance metric (reported relative to the AUPRC of a “null” model), although other measures of overall statistical performance (e.g., AUROC) are provided in supplementary information.

AUROC summarizes the tradeoff between true positives and false positives over a range of thresholds on the continuous model prediction score but is overly permissive in cases of imbalanced datasets–-although still suitable for assessing relative performance. The AUPRC metric summarizes the tradeoff between capturing all escape mutants (recall) and not incorrectly predicting escape mutants (precision). This approach is suitable for evaluating classification of imbalanced datasets but penalizes false positive predictions. In the case of escape predictors, false positive predictions may be due to insufficient sampling of the human antibody repertoire against the virus of interest, so this penalization is potentially too stringent. We normalize AUPRC by the “null” precision model AUPRC, which is equivalent to the fraction of escapes observed in the mutations experimentally screened. Therefore, AUPRC values are not comparable between viral proteins or subsets of DMS datasets with different fractions of escape mutations.

##### Comparison to known antibody footprints

We also evaluated the model’s ability to predict sites of antibody binding, as quantified by looking at antibody footprints in the RCSB PDB within a minimum all-atom distance of 3.5Å. Note that this is not information that is available to the model during training.

#### Comparison to pandemic data

##### Data Processing

We evaluate the model against occurrence of single mutations and strains in GISAID. In determining the set of Spike mutations to compare EVEscape scores to GISAID data, we consider only those mutations that are a single RNA nucleotide mutation distance from Wuhan. The date of lineage emergence is the 1^st^ percentile of dates for that variant (to avoid issues with outliers from GISAID data entry). Variants are marked as high frequency VOCs if their count is greater than 5,000 and it occurs in the first time period (pandemic divided into 12 periods) that any strain of that PANGO lineage appears. We define PANGO lineages for the VOCs by the nonsynonymous Spike consensus mutations for that strain from COVID-19 CG that occur in greater than 90% of strain sequences, ignoring insertions and deletions. Number of occurrences in the pandemic is defined by raw counts of GISAID records with a given substitution or set of substitutions.

##### Metrics

We calculate the fraction of predicted mutations (top 10%) seen in the pandemic over 100 times. We expect to see an increase in this fraction over the course of the pandemic, as more variants are observed and adaptive immune pressure increases with a growing vaccinated or previously infected population. We also calculate for each observed pandemic frequency minimum threshold, the percentage of pandemic mutants seen above that observed threshold that are predicted in the top 10%. We do not expect all pandemic mutants to be captured in the top 10% of predictions, because not all pandemic mutants are related to escape. Even amongst very frequent pandemic mutations mostly present in Variants of Concern, which we expect to be more enriched for high escape potential, we do not expect all of these mutations to be related to escape as some instead influence ACE2 binding or structural changes. To evaluate strain scores, we calculate the number of strains (and the corresponding percentile) that would need to be tested to have detected selected VOCs from all new strains in the two-week window they emerged. Unique new strains are defined by unique sets of Spike substitution mutations.

##### Escape within clinical antibody epitopes

We look at EVEscape predictions in the footprints (within 3.5Å) of six different clinical antibody epitopes. We then notate which of these mutations have already occurred in the pandemic (observed more than 10,000 times) and which have experimental evidence of escape for those clinical antibodies as seen in CoV-RDB^47^. We list all possible mutations, not just those a single nucleotide distance from Wuhan.

#### Comparison to strain neutralization

We show spearman correlation with experimental strain neutralization data as well as the linear regression line shown with a 95% confidence interval. EVEscape scores for these strains are calculated based on the mutations used in the experiment for each strain, ignoring indels. We convert percent reduction in neutralization (x) to fold reduction (1/1-x).

#### Regional Enrichment

We examine the distribution of EVEscape predictions throughout the Spike protein and, within the RBD, between the known footprints of different antibody classes^84^. We analyze enrichment of regions by comparing the average EVEscape score for the region to a distribution of the average EVEscape score of random regions. For comparison to full Spike, we compare to the scores of 500 random contiguous regions (of the same length as the region of interest) within Spike. For comparison to RBD, we compare to scores of 100 contiguous regions, using the full Spike model. We similarly compare scores of known neutralizing subregions to random regions in their respective full regions. We also compare enrichment of number of sites in the top 15% of EVEscape scores in each region relative to the length of the region. We consider the regions: NTD (sequence positions 14 - 306), RBD (319 - 542), S1* (543 - 685), and S2 (686 - 1273), where S1* refers to the region in S1 between RBD and S2. NTD and RBD are enriched in antibody sites. We also calculate the mutational tolerance of each region, the average EVE fitness score.

#### Epistasis

We analyze epistasis by comparing EVE scores on a Wuhan full Spike model (using a pre-pandemic alignment) and on an omicron (BA.2) full Spike model (using an alignment with data up to BA.2). The BA.2 epistatic shift is the Wuhan linear regression residual for a model fit to the two sets of EVE scores for all single mutations to full Spike. We compare the epistatic shift of two subsets of mutations, convergent omicron mutations and wastewater mutations^64^, to the full set of single mutations to full Spike. We also analyze the locations of the maximum epistatic shift, in relation to the Spike structure and to the set of sites mutated within BA.2.

#### Comparison to other computational models

We compare published SARS-CoV-2 RBD and Spike models predictions^20,22,52,85^ using metrics from above relevant to the intended purpose of each model (fitness or escape of either single mutations, sites, or strains).

## Footnotes

[1] Code and extended analyses are available at https://github.com/OATML-Markslab/EVEscape and at evescape.org

