## Supplementary Information for "Learning from pre-pandemic data to forecast viral escape"

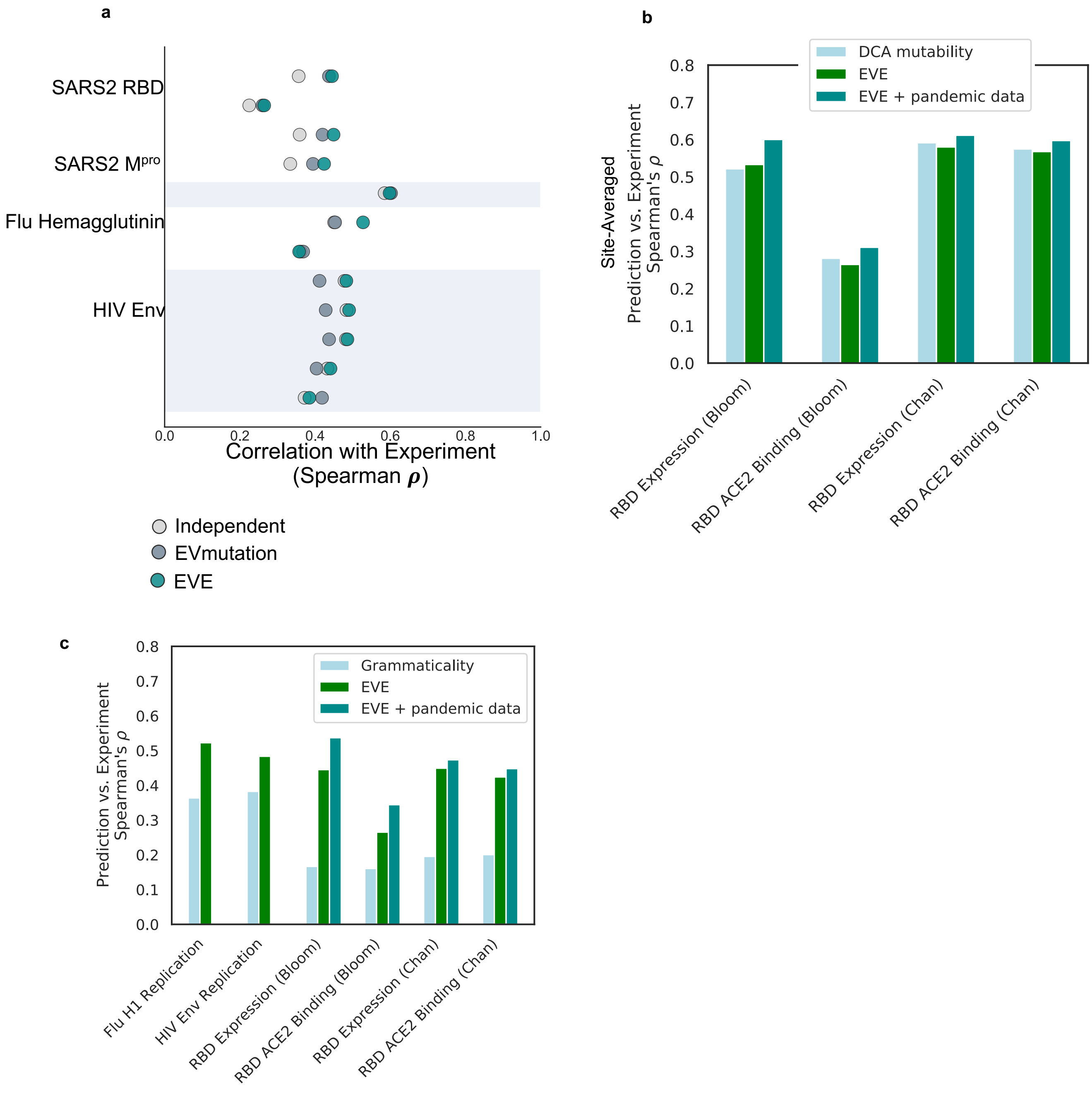


**Figure S1:Fitness effects of viral proteins predicted from evolutionary sequence models**. **a**) EVE predictions are well correlated with a broad range of viral surface protein deep mutation scanning experiments surveying protein replication and function, SARS-CoV-2 RBD[35,36] and $M^{pro}$[37], H1N1 hemagglutinin[31,32] and HIV env[30,33,34]. **b**) Site-averaged EVE predictions have similar correlations with site-averaged SARS-CoV-2 RBD DMS experiments as Potts model DCA[85] or EVmutation[24]. **c**) EVE predictions have higher correlations with Flu H1, HIV Env, and SARS-CoV-2 RBD DMS experiments than grammaticality in CSCS[52].

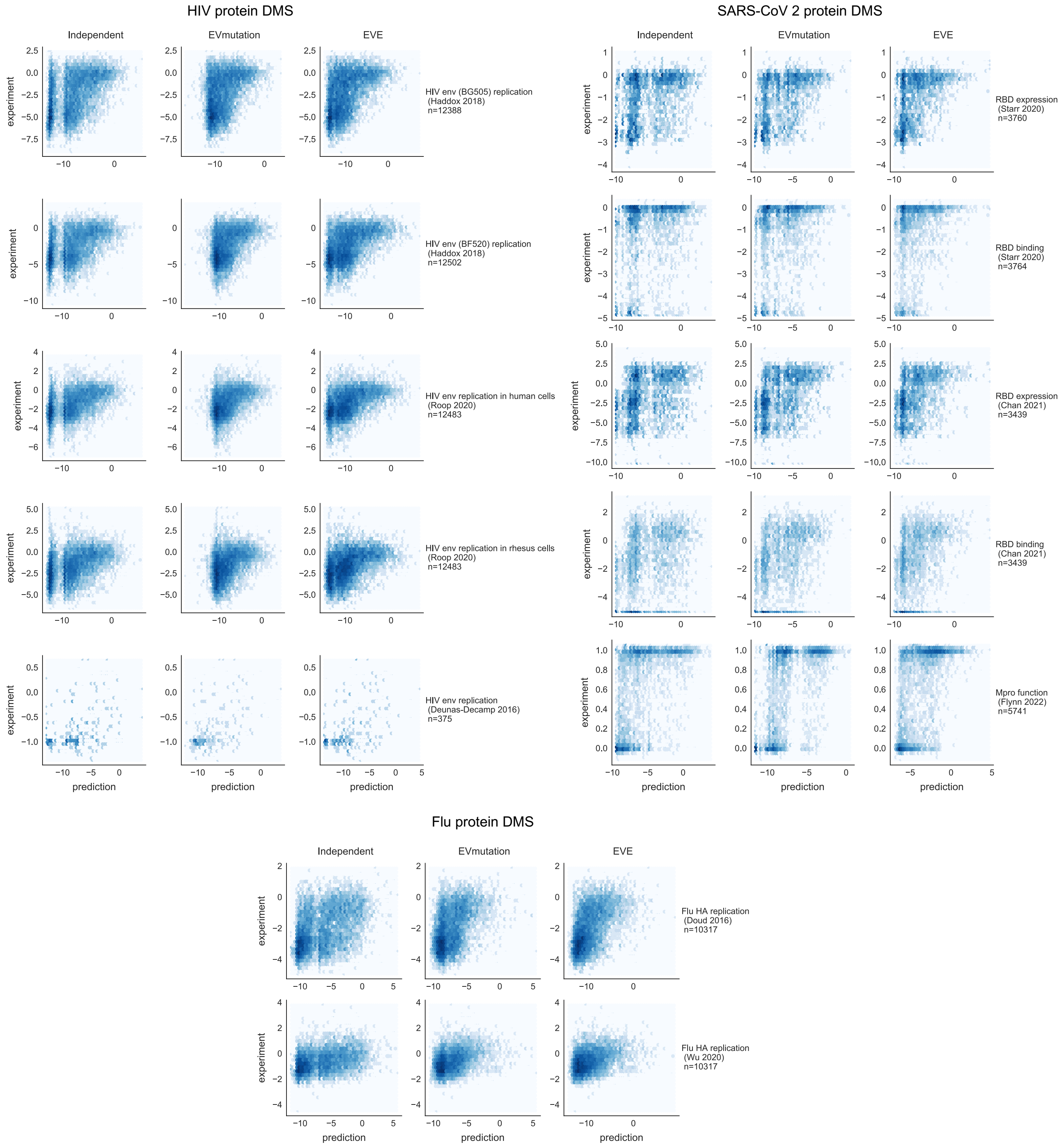


**Figure S2: Comparison of effects of mutations from experiment and computation.** Measurements of viral protein functions such as expression, replication and receptor binding in deep mutational scans versus predictions from an independent model, EVmutation, and EVE model.

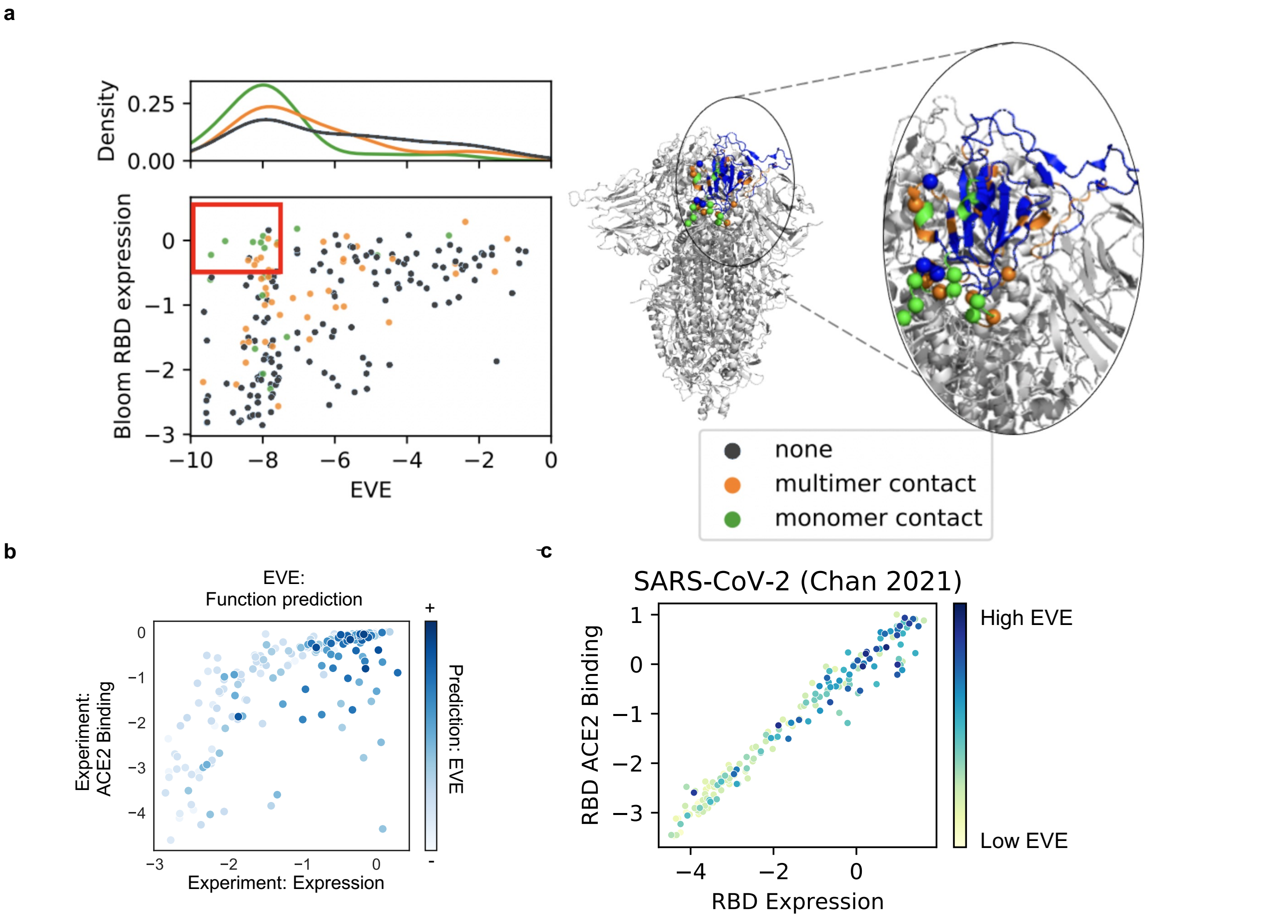


**Figure S3: EVE captures constraints beyond RBD expression assay. a**) Site-averaged EVE scores predict several sites that tolerate mutants in the yeast-display RBD expression assay[35] to be deleterious (red box)–many of these mutants are located at the interface between RBD and the rest of Spike protein. Sites in the red box in scatterplot are shown as spheres on the Spike structure (PDB: 7CAB). **b**) EVE prediction captures a combination of SARS-CoV-2 RBD yeast expression and ACE2 binding - features both necessary for successful immune escape (EVE spearman with expression = 0.45, EVE spearman with ACE2 binding = 0.38 when low expressed are removed)[35] **c**) The mammalian-cell RBD expression and ACE2 binding experiments are highly correlated, likely due to the alternate FACS-binning strategy and metric used for this ACE2 binding experiment[36]. EVE predictions are correlated with both measures.

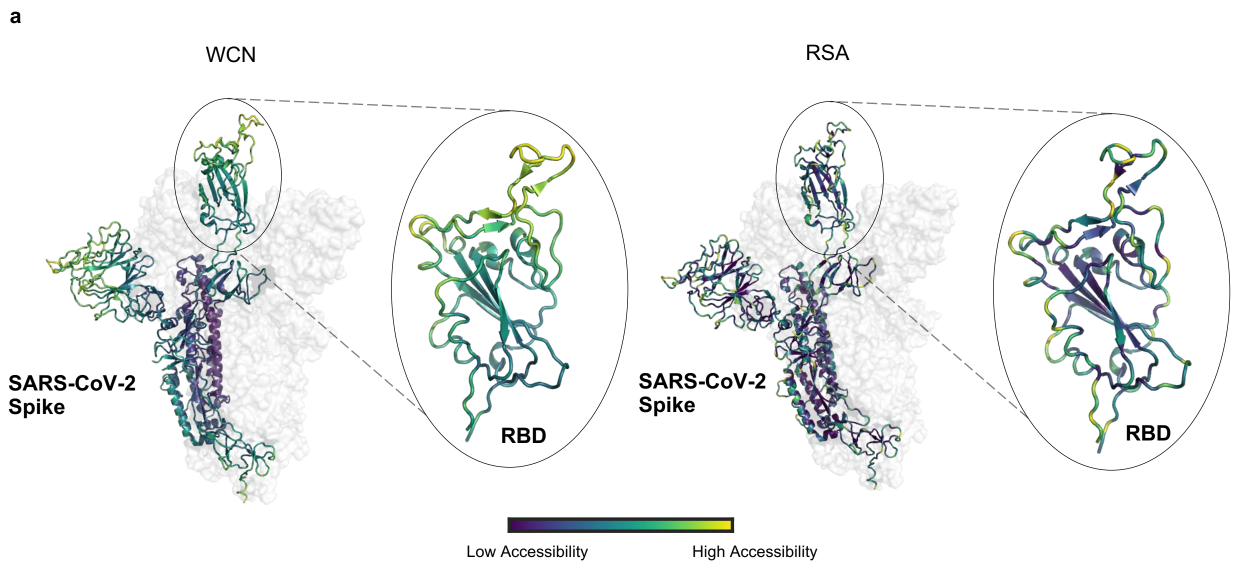


**Figure S4: Weighted contact number captures flexible regions. a**) WCN and RSA values visualized on the SARS-CoV-2 Spike structures show different distributions, particularly in the RBD (PDB: 7BNN), as WCN captures protrusion from the core structure.[38-41]

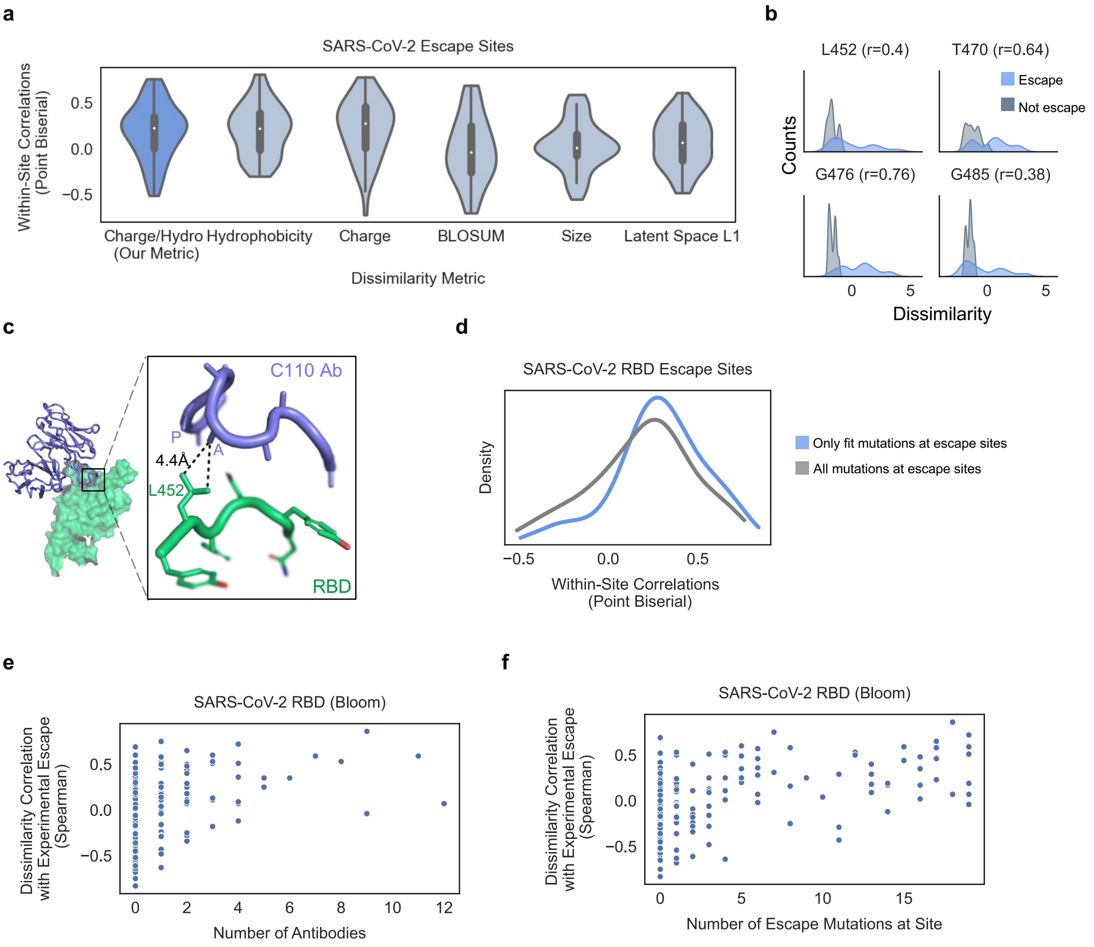


**Figure S5: Charge-hydrophobicity metric captures residue dissimilarity relevant for loss of antibody binding. a**) Within-site point biserial correlations between residue dissimilarity metrics and SARS-CoV-2 DMS escape data at escape sites (sites with 3-17 escape mutations). More sites have a higher correlation for our charge-hydrophobicity metric than charge or hydrophobicity alone, BLOSUM62, residue size, or EVE latent space (L1) distance. **b**) Charge-hydrophobicity dissimilarity performance in key sites **c**) The L452 RBD site is an example of decrease in hydrophobicity displacing the proximal alanine in the RBD C110 antibody interaction. (PDB: 7K8V) **d**) Within-site correlations at RBD escape sites increase when considering only mutations where fitness is maintained (passes Bloom lab's RBD expression and ACE2 binding cutoffs) **e**) Within-site correlations between residue dissimilarity and escape increase when more antibodies have escape mutations at that site. **f**) Within-site correlations between residue dissimilarity and escape increase when more mutations escape at site (and there can be no correlation with binarized escape when every mutation escapes).

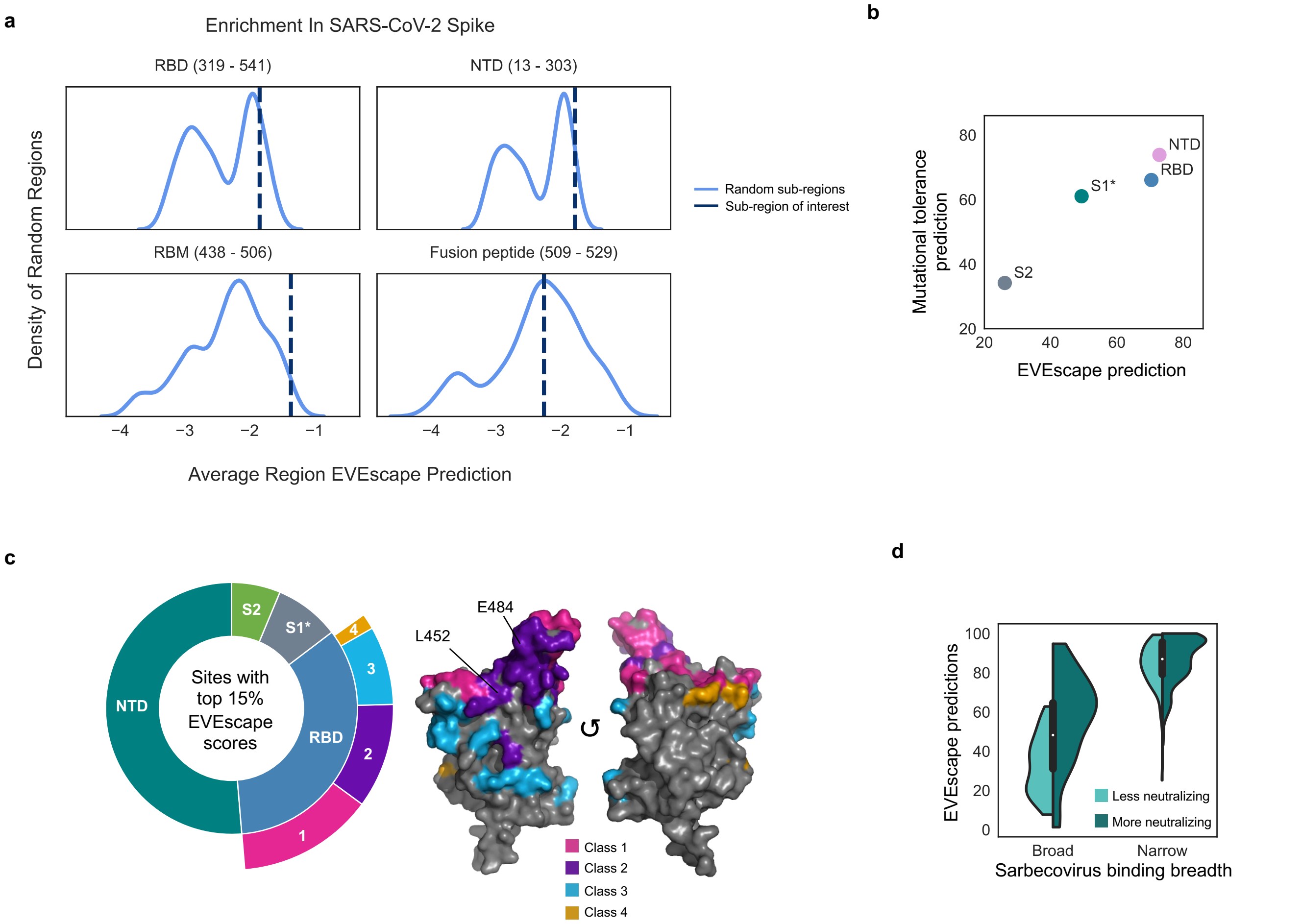


**Figure S6: EVEscape enrichment in regions of SARS-CoV-2 Spike. a**) RBD (particularly receptor binding motif (RBM)) and N-terminal domain (NTD) have significantly enriched average EVEscape scores, relative to a distribution of 500 random contiguous regions of the same length from full Spike. **b**) Average region EVEscape predictions are highest in RBD and NTD, though NTD is more mutationally tolerant with a higher average region EVE fitness score. **c)** EVEscape predictions cover diverse epitope regions across Spike and diverse RBD antibody classes[84] (3D structure of RBD on the right), including known immunodominant sites (E484, K417, L452) (PDB ID: 7BNN). The regions considered are NTD (sequence positions 14 - 306), RBD (319 - 542), S1* (543 - 685), and S2 (686 - 1273), where S1* refers to the region in S1 between RBD and S2. **d**) EVEscape scores experimental escape mutants from narrow antibodies and broad neutralizing antibodies higher than those from broad, non-neutralizing antibodies. Sarbecovirus binding breath and neutralization from Starr et al.[9]

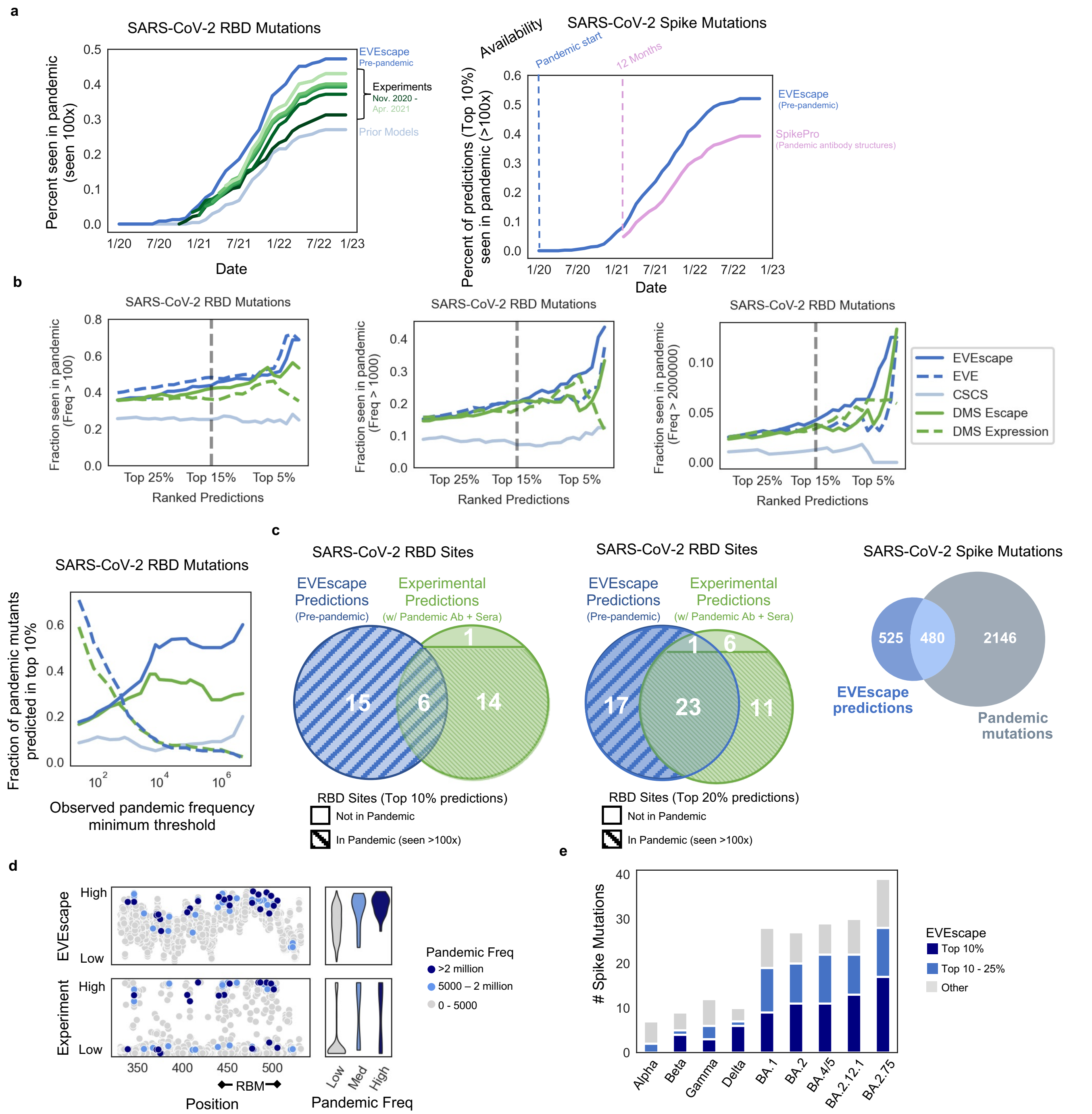


**Figure S7: EVEscape as accurate as experimental scans at anticipating pandemic variation: retrospective analysis. a**) Fraction of RBD predictions in top 15% of EVEscape, DMS experiments (Bloom Set, Table S4), and prior models[52] seen by each date over 100 times in GISAID (left). DMS experiments are separated into which studies were available by each starting date. EVEscape predictions for for full Spike and prior SpikePro model[21] (right). **b**) Fraction of mutations seen 100, 1000, or 2 million times over different thresholds of top ranked predictions (Top) and share of predicted escape mutations in top decile of prediction based on their observed frequency (Bottom). **c**) Venn diagram of RBD sites seen in the top 10% (left) or top 20% (middle) of EVEscape and all DMS experimental predictions (Bloom Set Table S4), with markings for whether the site was seen over 100 times in GISAID over the full pandemic.  Venn diagram of full Spike sites seen in top 10% of EVEscape and seen >100 times over the full pandemic (right). **d**) Comparison of  EVEscape computational model predictions (top panel, y axis EVEscape score) and DMS experimental predictions (bottom panel, y axis experimental score) to frequency of mutations. **e**) The majority of Spike mutations in VOC strains have high EVEscape scores.

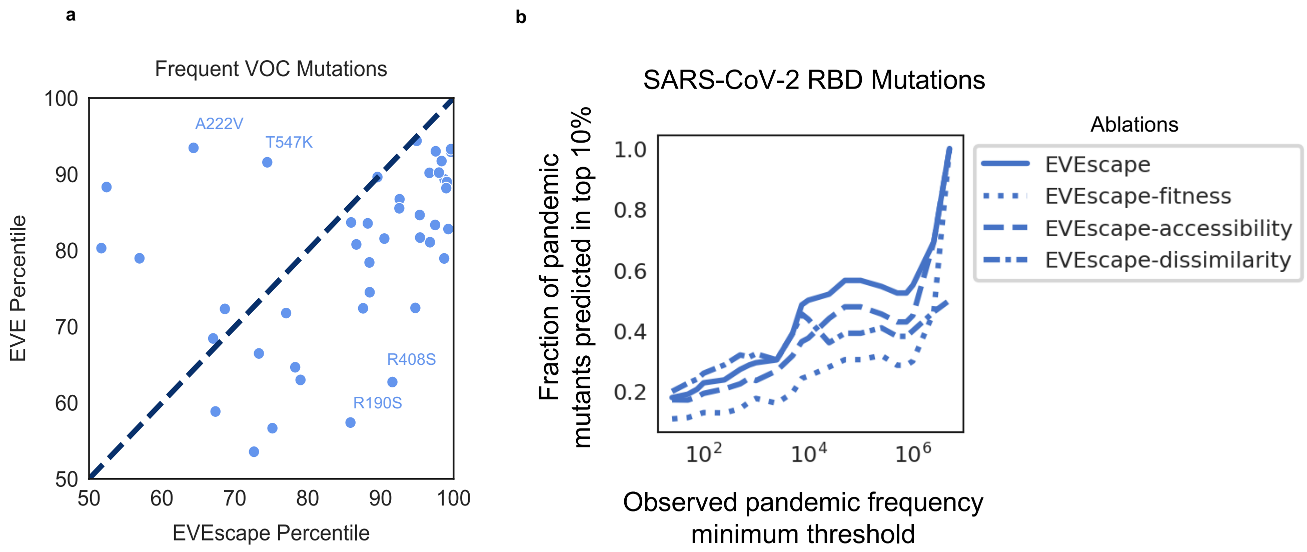


**Figure S8: The role of EVEscape components in capturing pandemic variant mutations**. **a**) EVEscape is more predictive than EVE alone at capturing frequent VOC mutations in full Spike. VOC mutations with high EVE scores and lower EVEscape scores (i.e., A222V and T547K) are known to impact structure and to not escape sera neutralization. Mutations with the highest EVEscape but low EVE scores (i.e., R190S and R408S) are in hydrophobic pockets that may promote antibody binding[54]. **b**) EVEscape is more predictive of high-frequency pandemic mutations than ablations of any of its 3 components. Notably, the ablation of the dissimilarity term leads to similar performance at identifying low-frequency mutations, but inferior performance at identifying high-frequency mutations.

| Clinical therapeutic antibody | Pre-pandemic forecasting of mutations in epitopes |
|---|---|
| Bamlanivimab | T470R, T470K, **S494R**, **Q493R**, **Q493L**, **Q493K**, **Q493H**, **G485R**, G485E, G482R, G482D, **E484V**, **E484Q**, **E484K**, E484G, **E484A** |
| Sotrovimab | K444T, K444N, K444M, K444I, K444E, G485R, G485E, G446R, G446E, G446D, E484V, **E484Q**, **E484K**, E484G, **E484A** |
| Etesevimab | S443R, **N440K**, L441R, L441H |
| Imdevimab | R346W, R346T, R346S, R346P, R346M, R346L, R346I, R346G, R346C, Q498R, Q498K, **N440K**, L441R, L441H, **K444T**, **K444N**, **K444M**, **K444I**, **K444E**, **G446R**, G446E, **G446D** |
| Casirivimab | **Q493R**, **Q493L**, **Q493K**, **Q493H**, L455R, K417M, K417I, **K417E**, G485R, G485E, **E484V**, **E484Q**, **E484K**, E484G, **E484A** |
| Regdanvimab | S494R, **Q493R**, Q493L, Q493K, Q493H, L455R, **L452R**, K417M, K417I, K417E, E484V, E484Q, **E484K**, E484G, **E484A** |

Pandemic mutations (Freq >10,000) colored & Experimental evidence **bolded**

**Figure S9: Forecasting of clinical antibody epitope escape mutations.** Forecasted mutations from the pre-pandemic model intersected with six clinical therapeutic monoclonal antibody epitopes. Epitopes are defined by sites within 3.5Å. Experimental evidence is from CoV-RDB[47]. All possible mutations are considered (not just those a nucleotide distance of one from Wuhan).

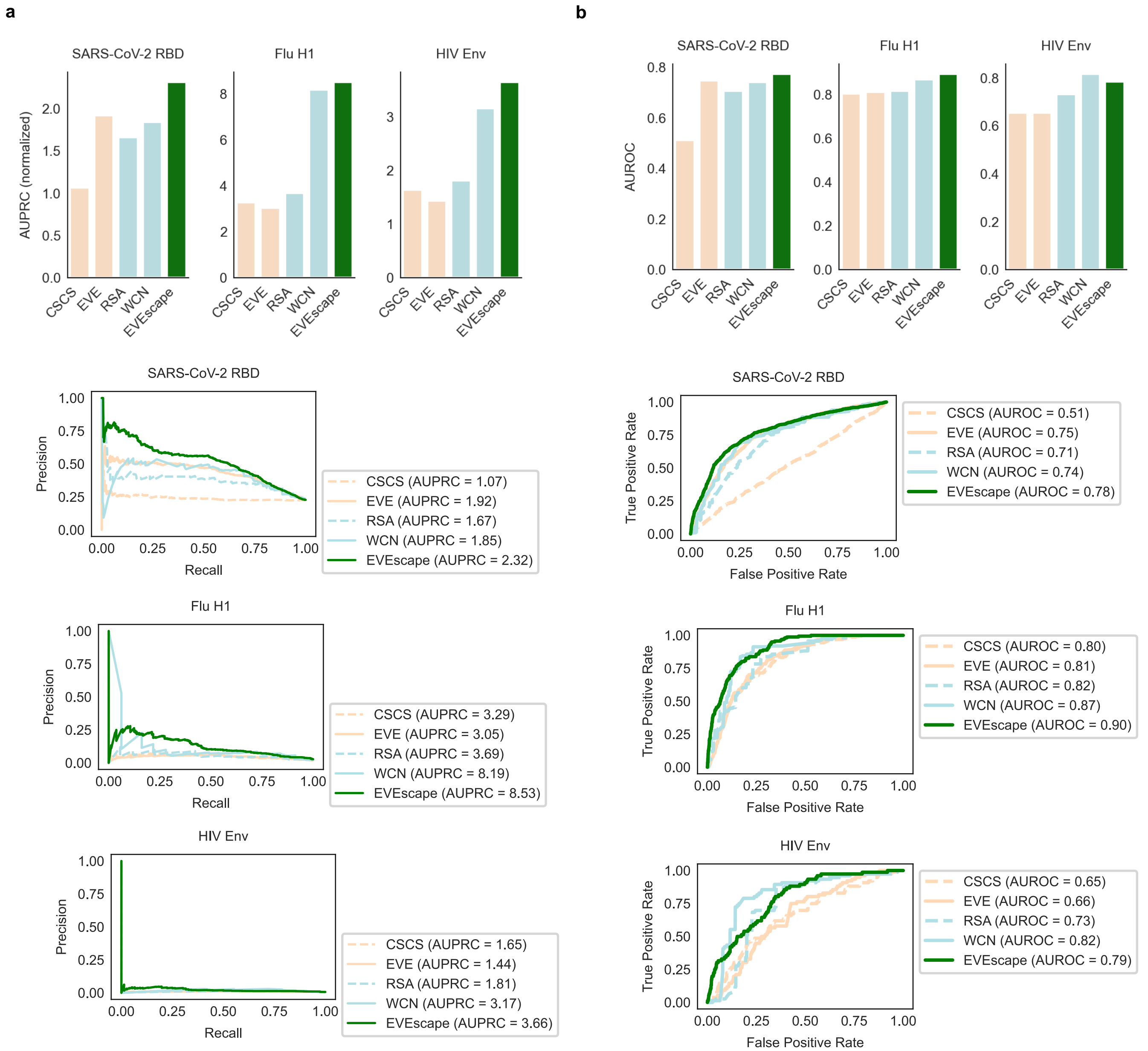


**Figure S10: EVEscape performance on escape DMS data is generalizable across viruses.** Precision-Recall (with AUPRC normalized by "null" model) (**a**) and AUROC (**b**) of predicting DMS escape mutations, for SARS-CoV-2 RBD, Flu H1, and HIV Env.

Note: The "null" model AUPRC is equivalent to the fraction of observed escapes, and therefore AUPRC values are not comparable between viral proteins with different fractions of escape mutations (i.e. RBD and HIV Env). The fraction of observed escapes in the DMS experiments are 0.19 for RBD, for 0.015 for Flu, and 0.006 for HIV – Flu and HIV data examined far fewer antibody and sera samples (Table S5).

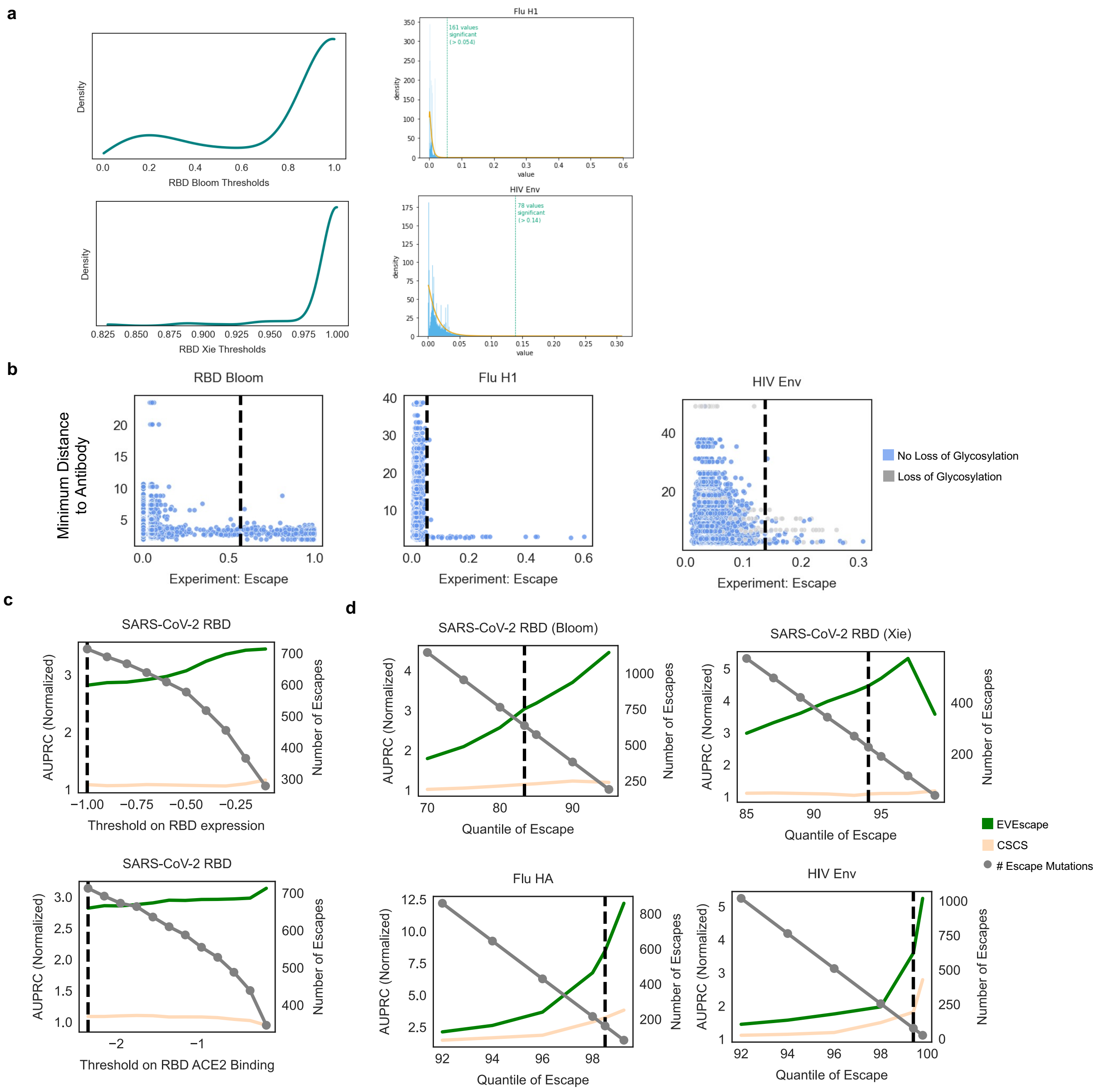


**Figure S11: EVEscape performance is robust across data thresholds. a**) Distribution of escape thresholds from bootstrapping 8 antibodies 1000 times and fitting a gamma distribution to each sample for Bloom and Xie RBD escape data (left) and gamma distributions to select Flu and HIV escape thresholds (right). **b**) Maximum escape values (over set of antibodies with PDB structures) for each mutation vs. the minimum distance to an antibody—most escape mutations (to the right of dashed line) are to residues with atoms  to within 5Å of any residue on the antibody. For HIV, this is true for the mutations that do not involve loss of glycosylation. **c**) Impact of choice of RBD expression and ACE2 binding thresholds (dashed line uses thresholds chosen by Bloom escape papers and our paper) on AUPRC (normalized by “null” model – fraction of observed escapes) and # of mutations considered as escape. d) Impact of choice of escape threshold on RBD (Bloom and Xie data separated), Flu, and HIV AUPRC (normalized) and # of escape mutations (dashed line uses escape threshold chosen by our paper).

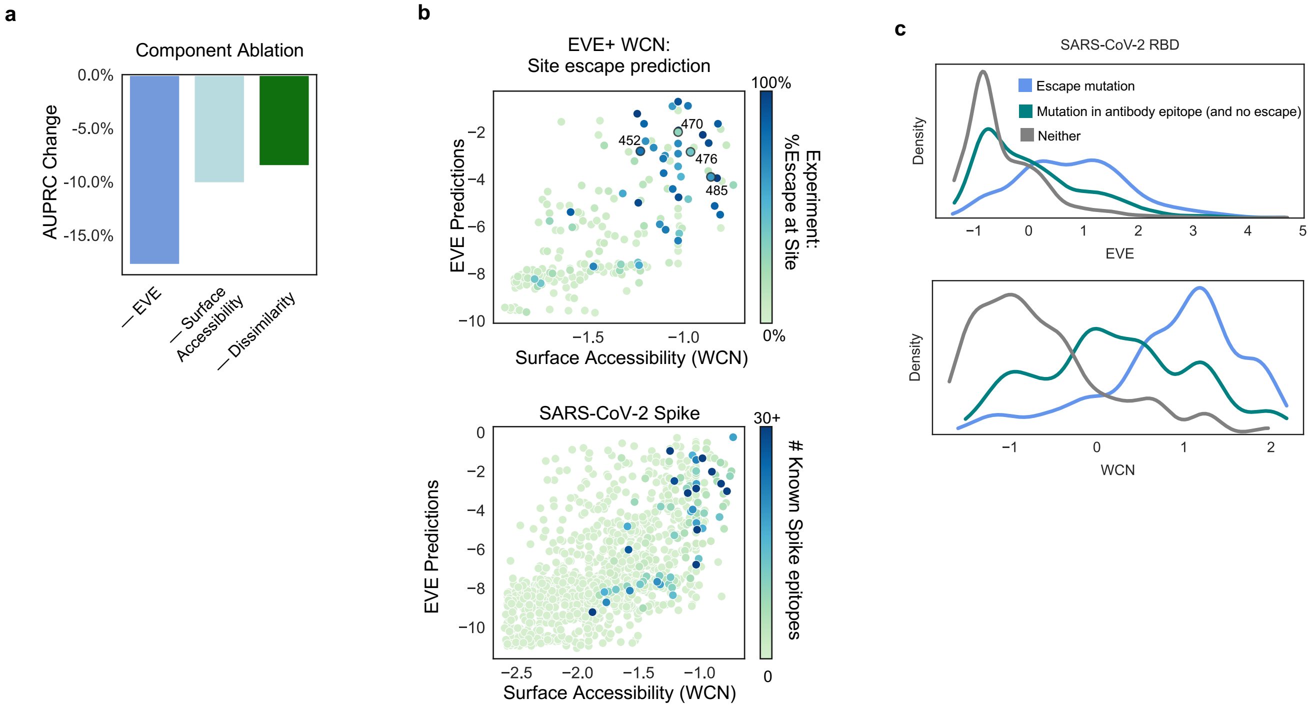


**Figure S12: Surface accessibility metrics, mutation effect models, and dissimilarity provide complementary information for predicting antibody epitopes and escape mutations. a)** All features of EVEscape contribute to performance in predicting RBD escape mutants. **b**) Sites with either high WCN accessibility or high EVE fitness predictions have a greater percent of escape mutants (upper). WCN and EVE predictions provide similar information about the location of Spike epitopes as represented in antibody-Spike crystal structures in RCSB PDB (lower). **c)** Density of standard-scaled EVEscape components differ for SARS-CoV-2 RBD escape (and antibody epitopes) and non-escape mutations.

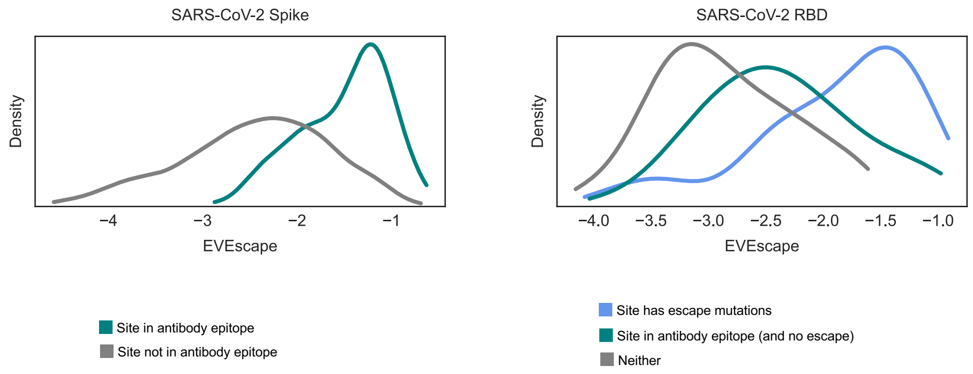


**Figure S13: Top EVEscape predicted sites are known escape and in antibody footprints.** Density of site-averaged EVEscape for SARS-CoV-2 full Spike (left) and RBD (right) shows success of EVEscape at distinguishing sites with observed escape mutations, as well as sites in known antibody epitopes, from sites with no evidence of antibody binding or escape. All but 2 sites in the top 20% of EVEscape scores are in known antibody footprints or have escape mutations in experiments.

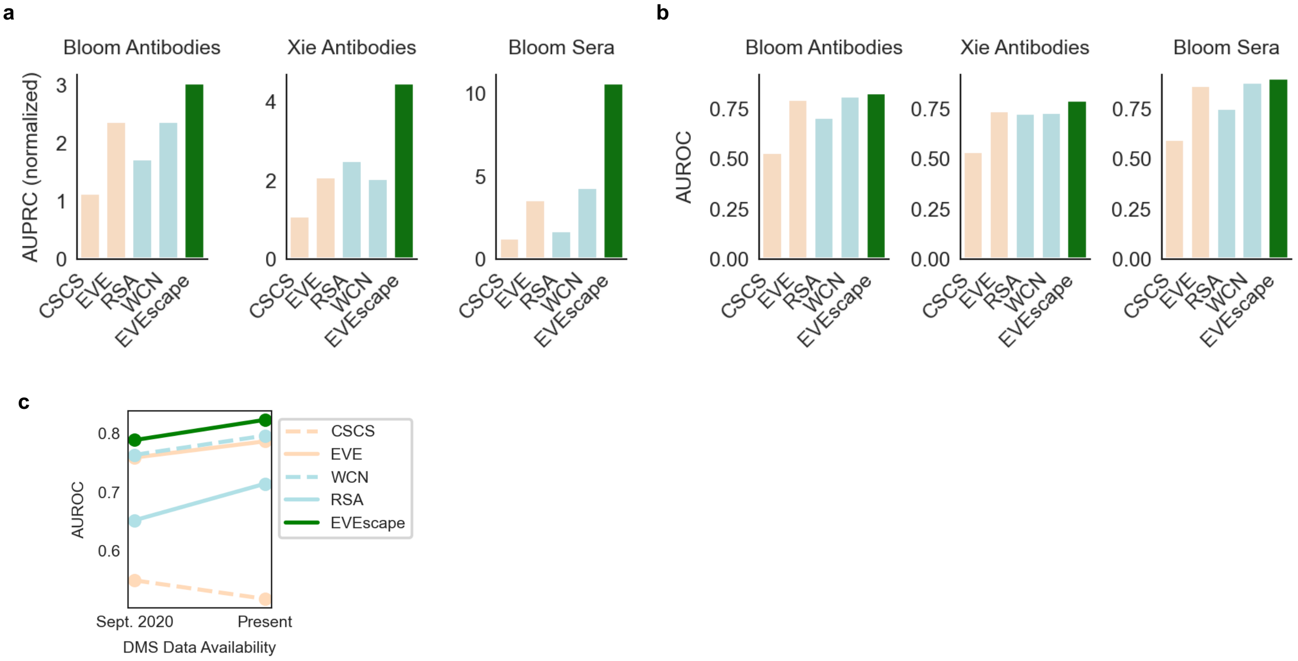


**Figure S14: EVEscape RBD performance is robust to antibody and sera samples and improves with more available data for validation.** Precision-Recall (with AUPRC normalized by "null" model) (**a**) and AUROC (**b**) of predicting RBD DMS escape mutations, for Bloom and Xie antibodies and Bloom sera. **c**) Comparison of model performance (AUROC) between data from first escape DMS study (10 antibodies – Sept. 2020)[4] and data available at present (338 antibodies, 55 sera samples).
Note: The "null" model AUPRC is equivalent to the fraction of observed escapes, and therefore AUPRC values are not comparable between data samples with different fractions of escape mutations (i.e., Bloom sera vs. Bloom antibodies, Table S5). The fraction of observed escapes in the DMS experiments are 0.17 for Bloom Ab, 0.06 for Xie Ab, and 0.003 for Bloom sera.

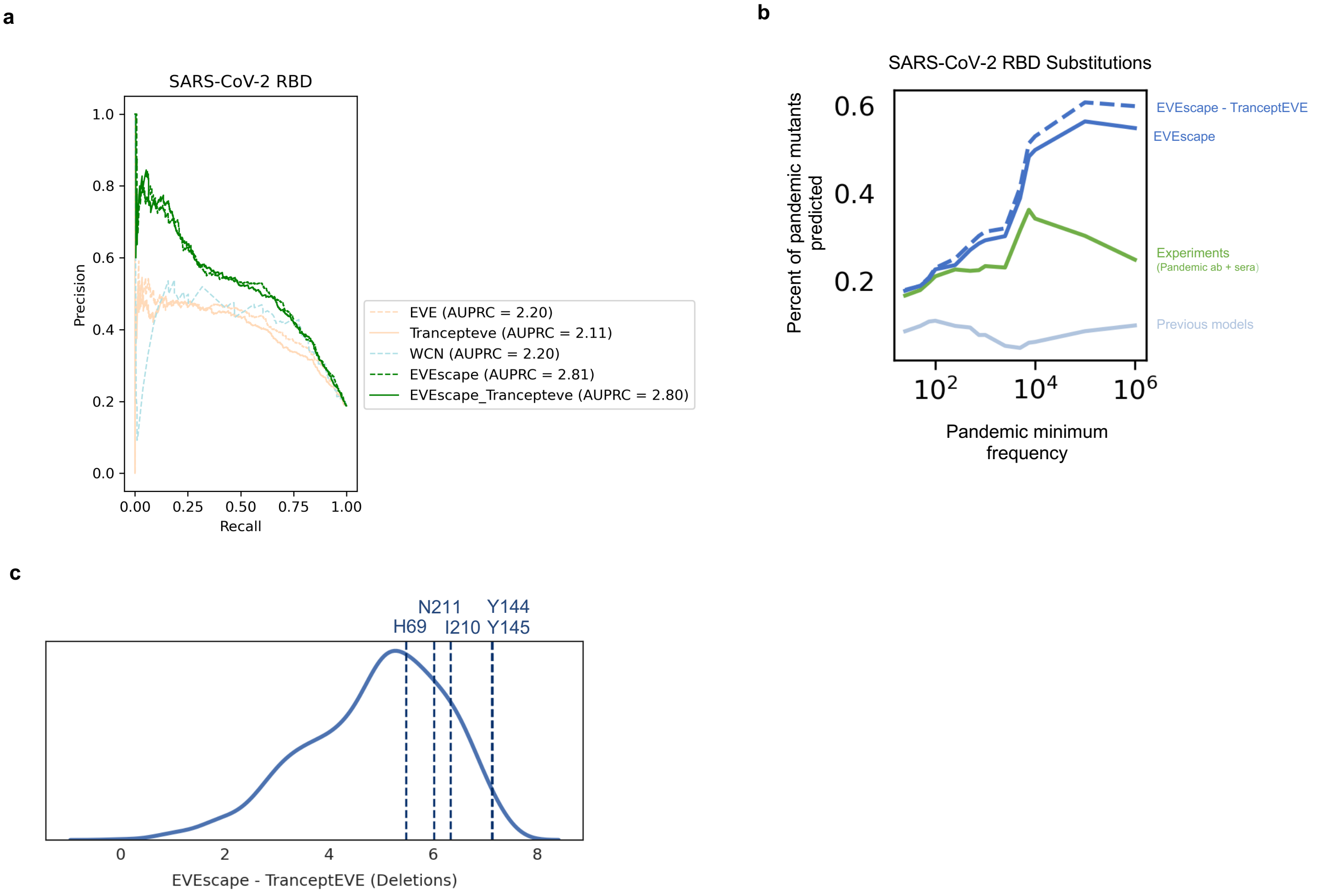


**Figure S15: EVEscape adapts to new transformer model of mutation fitness capable of scoring indels.** The EVEscape fitness component can be substituted with a new generative model, Trancept-EVE[59] that is capable of scoring substitutions as well as insertions and deletions. **a)** EVEscape using TranceptEVE as the fitness model performs equivalently to EVEscape using EVE at predicting substitutions from from deep mutational scans that escape antibody binding. **b)** Percent of predicted substitutions in top decile of prediction based on their observed frequency during the pandemic shows EVEscape with TranceptEVE is just as good as, or better than, EVEscape using EVE at predicting pandemic substitutions. **c)** Histogram of EVEscape scores with TranceptEVE as a fitness model for all single deletions to Spike. Single deletions seen in the pandemic more than 1000 times are predicted higher than most other single deletions, especially the very frequent pandemic deletion Y144- (seen more than a million times).

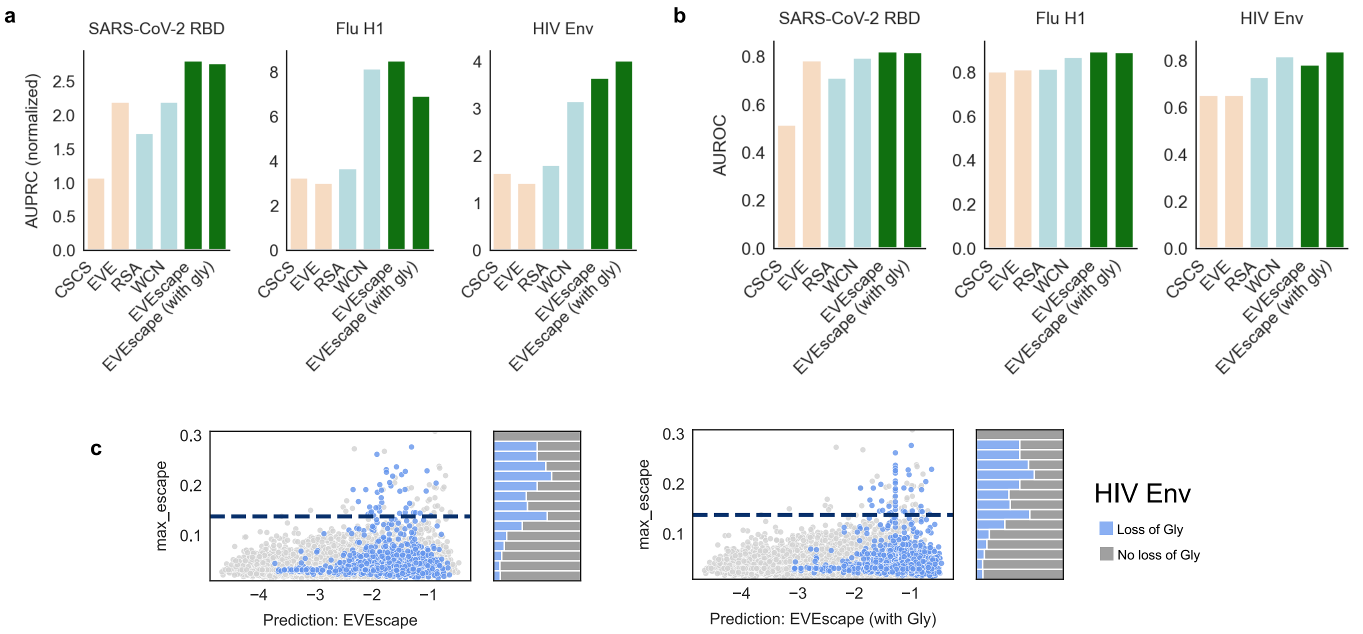


**Figure S16: Incorporating glycosylation in EVEscape improves performance on HIV Env.** Precision-Recall (with AUPRC normalized by "null" model – fraction of observed escapes) (**a**) and AUROC (**b**) of EVEscape and EVEscape+Gly predicting DMS escape mutations for SARS-CoV-2 RBD, Flu H1, and HIV Env. **c**) Scatterplot of HIV Env maximum escape at each mutation vs. EVEscape predictions with and without glycosylation. Hue indicates mutations that cause cause loss of glycosylation. The majority of HIV Env escape mutations involve glycosylation loss, and EVEscape+Gly performs better on these mutations.

Note: In the limited HIV Env dataset examining 8 antibodies, 50% of all escape mutations are likely due to removal of a glycan[17]. The effects of glycosylation changes may not be reflected in the SARS-CoV-2 Spike experiments as these experiments were conducted in a yeast system with different surface glycan types[8]. While SARS-CoV-2 Spike (22 glycosylation sites) and Flu H1 (up to 11 glycosylation sites) are much less extensively glycosylated than HIV Env (up to 30 glycosylation sites), some glycosylation changes in these proteins facilitate escape[60-63].

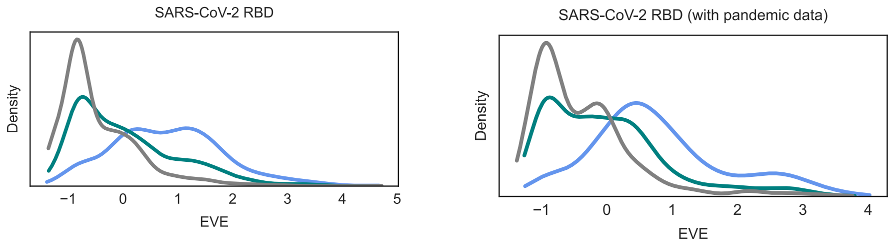


**Figure S17: Incorporating pandemic data into EVE improves prediction of escape DMS.** Incorporating pandemic sequences in EVE training data results in a greater distinction between escape and non-escape mutations with high EVE scores.

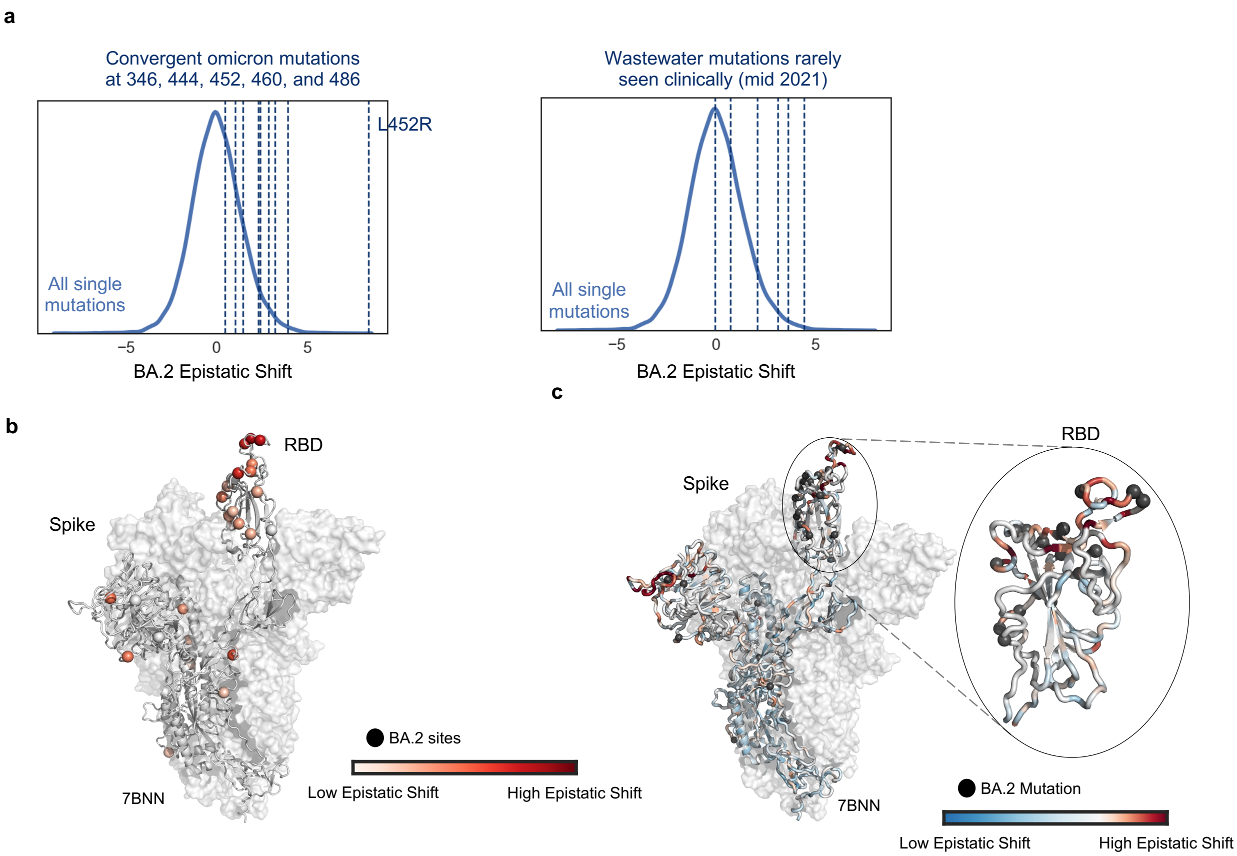


**Figure S18: EVEscape captures the epistatic shift between Wuhan and BA.2.**
**a**) Histogram of epistatic shift values between Wuhan and BA.2 EVE models for all single mutations, calculated as linear regression residuals. Convergent mutations that arise multiple times in Omicron lineages (mutations at sites 346, 444, 452, 460, and 486) highlighted on the left. Wastewater mutations seen mid-2021[64] that were rarely seen clinically in patients, and so likely epistatic (right). **b**) Max epistatic shift magnitudes of mutations at sites in BA.2 shows high epistatic shifts concentrated in RBD. **c**) Large epistatic shifts for mutations on Wuhan and BA.2 strains concentrated at sites proximal to BA.2 mutations.

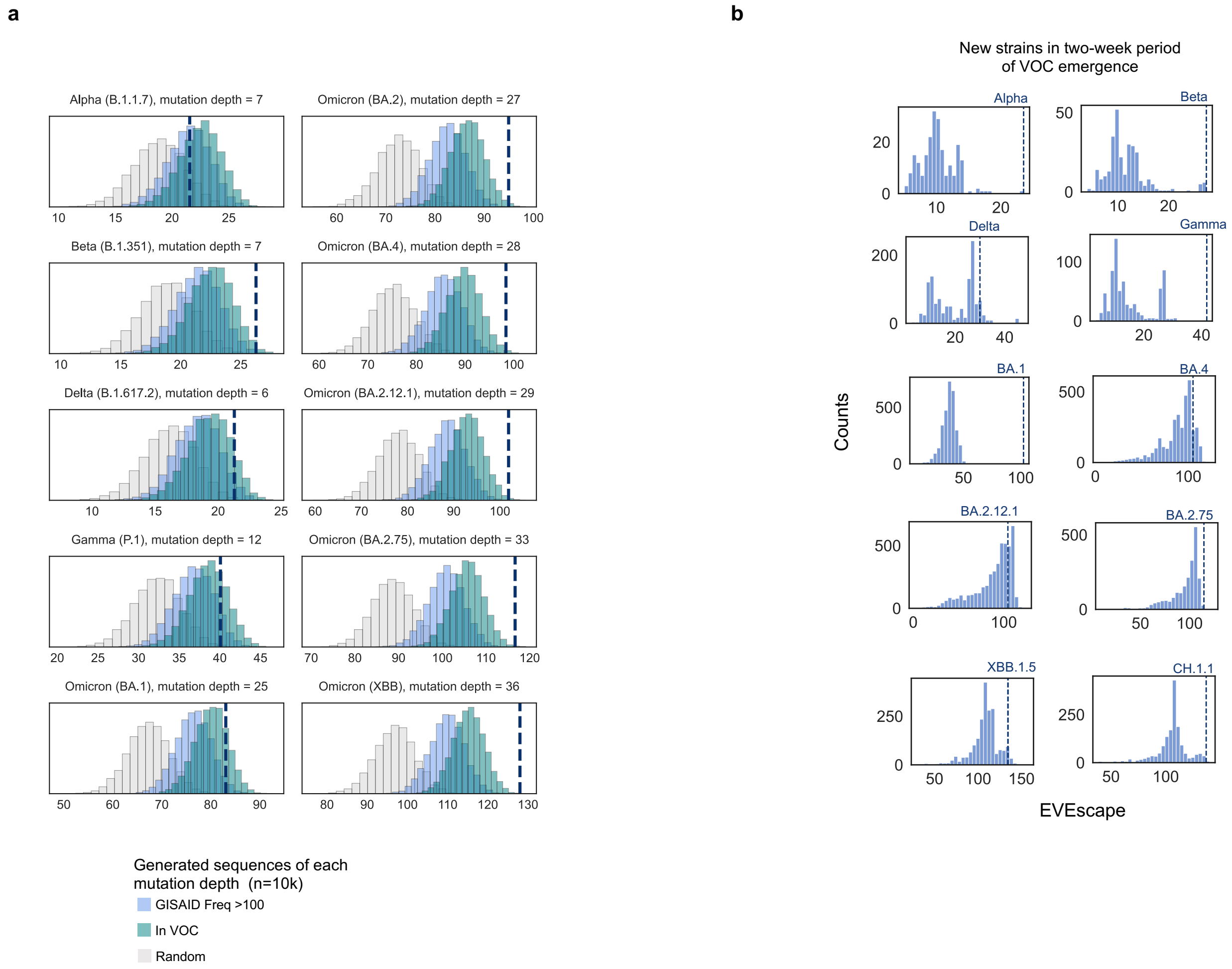


**Figure S19: EVEscape strains**

**a)** VOCs have high EVEscape scores compared to random mutations at the same mutation depth, particularly Beta and later Omicron strains**. b)** VOCs are among the highest scoring new strains for their two-week period of emergence using a pre-pandemic EVEscape model.

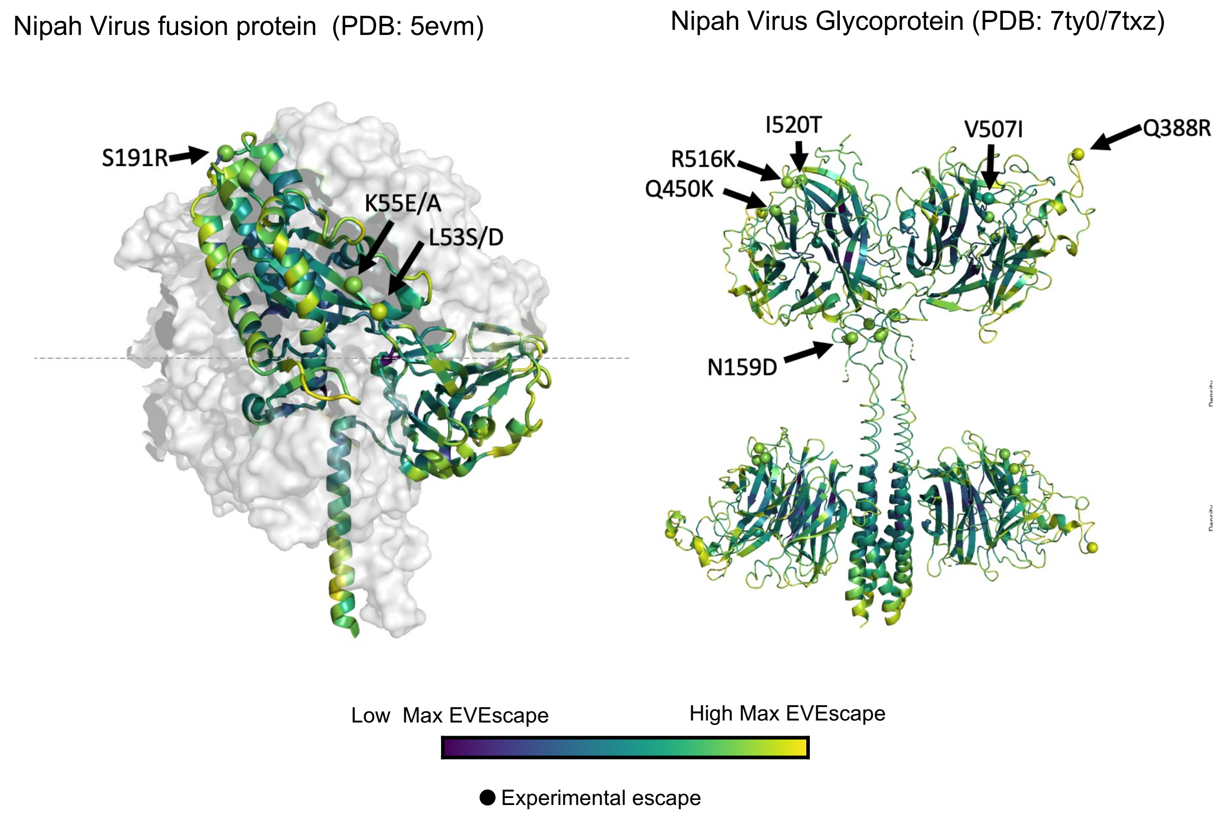


**Figure S20: EVEscape predictions for potential pandemics** Site-maximum EVEscape scores on Nipah Virus fusion protein (left) and Glycoprotein (right) structures depict regions of high EVEscape scores and known escape mutations with experimental evidence[68-72] (little is known for this understudied virus with pandemic potential) are highlighted with spheres.

| | RBD (+ pandemic data) | **RBD** | Spike (+ pandemic data) | **Spike** |
|---|---|---|---|---|
| SARS-CoV-2 | 1394 | 1 | 1292 | 1 |
| SARS-CoV-1 | 37 | 24 | 27 | 23 |
| Other SARS-like | 110 | 101 | 108 | 99 |
| MERS | 305 | 265 | 293 | 259 |
| Betacoronavirus 1 (OC43) | 577 | 416 | 504 | 394 |
| Alphacoronavirus 1 | 0 | 0 | 240 | 175 |
| 229E | 0 | 0 | 131 | 95 |
| NL63 | 0 | 0 | 52 | 47 |
| HKU1 | 57 | 27 | 54 | 27 |
| HKU15 | 0 | 0 | 212 | 141 |
| Avian coronavirus | 0 | 0 | 668 | 581 |
| Porcine epidemic diarrhea virus | 0 | 0 | 1800 | 1440 |
| Other coronavirus | 252 | 175 | 486 | 347 |
| Other/unknown | 0 | 0 | 1 | 0 |
| **Total** | **2732** | **1009** | **5868** | **3629** |

**Table S1: Taxa of sequences in Spike and RBD training alignments. RBD and Spike without pandemic data are the primary alignments used throughout this paper.**

| Protein | Prefix | # Sequences | Sequence Length | % Coverage | N_eff |
|---|---|---|---|---|---|
| Flu H1 | I4EPC4_t0.99_b0.1 | 71463 | 565 | 97.0% | 12238.5 |
| HIV Env | Q2N0S5_20-709_b0.1_t0.99 | 109050 | 690 | 97.2% | 48082.1 |
| SARS-CoV-2 RBD (+ pandemic data) | P0DTC2_321-541_b0.3 | 2732 | 221 | 98.6% | 277.7 |
| SARS-CoV-2 RBD | P0DTC2_321-541_b0.3_sc0.5_cc0.3_pre2020 | 1009 | 221 | 98.6% | 195.4 |
| SARS-CoV-2 Spike (+ pandemic data) | P0DTC2_sc0.5_cc0.3_b0.1 | 5868 | 1273 | 98.9% | 1639.1 |
| SARS-CoV-2 Spike | P0DTC2_sc0.5_cc0.3_b0.1_pre 2020 | 3629 | 1273 | 94.7% | 1345.6 |
| Lassa Glycoprotein | GLYC_LASSJ_deep_b0.05 | 1093 | 491 | 99.8% | 536.2 |
| Nipah Glycoprotein | GLYCP_NIPAV_b0.05 | 5036 | 602 | 88.7% | 1328.1 |
| Nipah Fusion Protein | FUS_NIPAV_b0.05 | 6155 | 546 | 94.1% | 1105.6 |

**Table S2: EVE training alignment summary statistics.**

| Virus | Protein | Study | Strain | Alignment | Assay variable | N | $\rho_{Independent}$ | $\rho_{EVmutation}$ | $\rho_{EVE}$ |
|---|---|---|---|---|---|---|---|---|---|
| Influenza | H1 | Doud 2016[31] | A/WSN/1933 | A0A2Z5U3Z0_9INFA_b0.1 | replication | 10317 | 0.45 | 0.45 | **0.53** |
| | | Wu 2020[32] | H1 (strain) | A0A6H1V8E8_9PLVG_Y373S_b0.1 | replication | 10317 | 0.36 | **0.37** | 0.36 |
| HIV | Env | Haddox 2018[30] | BG505 | A0A192B1T2_9HIV1_b0.1 | replication | 12388 | **0.48** | 0.41 | **0.48** |
| | | | BF520 | ENV_HV1B9_S364P-M373R_b0.1 | replication | 12502 | 0.48 | 0.43 | **0.49** |
| | | Roop 2020[33] | BG505 | A0A192B1T2_9HIV1_b0.1 | replication (human cells) | 12483 | 0.48 | 0.44 | **0.49** |
| | | | | | replication (rhesus cells) | 12483 | 0.43 | 0.40 | **0.44** |
| | | Duenas-Decamp 2016[34] | BG505 | A0A192B1T2_9HIV1_b0.1 | replication | 375 | 0.37 | **0.42** | 0.38 |
| SARS-CoV-2 | Spike RBD | Starr 2020[35] | Wuhan-Hu-1 | P0DTC2_321-541_b0.3_pre2020.a2m | yeast expression (RBD) | 3798 | 0.36 | 0.33 | **0.45** |
| | | | | | ACE2 binding | 3802 | 0.23 | 0.16 | **0.26** |
| | | Chan 2021[36] | | | human cell expression (full Spike) | 3458 | 0.33 | 0.32 | **0.45** |
| | | | | | ACE2 binding | 3458 | 0.31 | 0.30 | **0.42** |
| | $M^{pro}$ | Flynn 2022[37] | Wuhan-Hu-1 | nsp5-YP_009725301_b0.1.a2m | yeast growth | 5741 | 0.58 | **0.60** | **0.60** |

**Table S3: Experimental details and EVE, EVmutation, and independent model performance (spearman correlations) for DMS fitness experiments.**

| | PDB ID | Description |
|---|---|---|
| SARS-CoV-2 Spike | 6VXX | Spike (closed state) |
| | 6VYB | Spike (open state) |
| | 7CAB | Spike (closed state with higher sequence coverage) |
| | 7BNN | Spike (open state with higher sequence coverage) |
| Flu H1 | 1RVX | 1934 H1 Hemagglutinin (similar to Bloom DMS sequence) |
| HIV Env | 5FYL | BG505 SOSIP.664 Env (prefusion) Trimer structure created using structural symmetry in Pymol (adapted from Dingens et al.)[17] |
| | 7TFO | BG505 SOSIP.664 Env (CD4-bound open state) |
| Lassa Glycoprotein | 7PUY | Lassa Virus Josiah Strain Glycoprotein |
| Nipah Fusion Protein | 5EVM | Nipah Fusion Protein (prefusion) |
| Nipah Glycoprotein | 7TY0 | Nipah Glycoprotein Malaysian Strain |
| | 7TXZ | Nipah Glycoprotein Malaysian Strain |

**Table S4: PDB structures capturing diverse protein conformations used for accessibility calculations.**

| | Papers | Assay Details | # of Mutations | # of Escape Mutations (using thresholds from our paper) | # of Antibodies/ Sera | Alignment |
|---|---|---|---|---|---|---|
| SARS-CoV-2 RBD (Wuhan-Hu-1) | **Bloom Lab (antibodies):** Dong 2021[3] Greaney 2021[4] Greaney 2021[5] Greaney 2021[7] Starr 2021[8] Starr 2021[9] Tortorici 2021[11] Starr 2021[10] | FACS-based yeast display screening of antibody binding | 3819 | 635 | 36 | P0DTC2_321-541_b0.3_pre2020.a2m |
| | **Bloom Lab (sera)**: Greaney 2021[5] Greaney 2021[6] Greaney 2021[7] | FACS-based yeast display screening of sera binding | 3819 | 15 | 55 | P0DTC2_321-541_b0.3_pre2020.a2m |
| | **Xie Lab**: Cao 2022[12] | MACS-based yeast display screening of antibody binding | 3819 | 227 | 247 | P0DTC2_321-541_b0.3_pre2020.a2m |
| Flu H1 (A/WSN/1933) | Doud 2018[16] | Screening viral cell entry in the presence of antibodies | 10735 | 161 | 6 | I4EPC4_t0.99_b0.1.a2m |
| HIV Env (BG505) | Dingens 2019[17] | Screening viral cell entry in the presence of antibodies | 12730 | 76 | 8 | Q2N0S5_20-709_b0.1_t0.99.a2m |

**Table S5: Escape DMS data used for EVEscape validation.**
